# Coordinated changes in thymic stromal and hematopoietic cells that define the perinatal to juvenile transition

**DOI:** 10.1101/2025.03.04.641047

**Authors:** Anusha Vasudev, Colin R. Moore, Aparna Calindi, Seung Woo Kang, Bryan R. Helm, Jayashree Srinivasan, Siddhartha Shah, Ruiting Zong, Nandini Singarapu, Scott Casey, Andrew N. Macintyre, Ken S. Lau, Laura P. Hale, Qi Liu, Nancy R. Manley, Lauren I.R. Ehrlich, Ellen R. Richie

**Affiliations:** Department of Epigenetics and Molecular Carcinogenesis, The University of Texas MD Anderson Cancer Center, Houston, TX 77030; Department of Molecular Biosciences, The University of Texas at Austin, Austin, TX 78712; Department of Genetics, The University of Georgia, Athens, GA 30602; Department of Biostatistics, Vanderbilt University Medical Center, Nashville, TN 37232; Epithlielial Biology Center and Department of Cell and Developmental Biology, Vanderbilt University School of Medicine, Nashville, TN 37232; Department of Pathology and the Human Vaccine Institute, Duke University School of Medicine, Durham, NC 27710

**Keywords:** perinatal thymus, perinatal thymus stromal cell atlas, perinatal thymocytes, perinatal thymic epithelial cells, perinatal thymic mesenchyme, perinatal thymic dendritic cells, perinatal thymocytes, IGF2, Interferon I/III

## Abstract

T cells in the perinatal thymus have distinct phenotypes and functions that may be instructed by age-specific features of the microenvironment. We evaluated molecular and cellular profiles of thymic stromal cells, including thymic epithelial cells (TECs), mesenchyme, endothelium, and hematopoietic antigen presenting cells (hAPCs), from birth through one-month of age in mice. Single-cell transcriptional profiling, flow cytometry, and immunohistochemistry revealed coordinated stromal changes accompanied by altered thymocyte differentiation at defined transitional ages during the shift from perinatal growth to juvenile homeostasis, which was mirrored in humans. These analyses link diminished IGF2 expression by mesenchymal cells with activation of the RB pathway in TECs at the transition. Moreover, a coordinated increase in type I interferon signaling in stroma across the transition is associated with altered antigen processing and presentation signatures in TECs and hAPCs. Collectively, these datasets provide a resource to interrogate thymic stroma across the perinatal to juvenile transition.

## INTRODUCTION

Unlike most organs in which postnatal growth is coordinated with increasing size of the organism, thymus growth is untethered to body size. Thymus weight and cellularity increase exponentially during the early perinatal period, but growth tapers off, and thymus size then begins to decline at a juvenile age^1–3^. The thymus plays a critical role in adaptive immunity, as it constitutes a unique microenvironment required for T cell development and selection^4,5^. Changes in thymic size across the lifespan directly correlate with the production and output of new T cells that contribute to immune responses^6^. Beyond changes in the number of T cells produced, growing evidence indicates differences in the composition and function of T cells generated in the perinatal versus juvenile or adult thymus^7,8^.

Multiple lines of evidence demonstrate that perinatal T cells have profoundly different functional potential compared to their adult counterparts. Mouse and human perinatal T cells are hyporesponsive to TCR stimulation; perinatal CD4+ T cells are more biased towards Th2 responses; and CD8 T cells mount short-lived, innate-like responses associated with elevated cytokine production and diminished memory formation^7–9^. Moreover, unique subsets of innate γδT cells that seed different organs are produced in waves in the fetal thymus^10,11^. Furthermore, cells produced in the perinatal thymus, such as virtual memory CD8 T cells and regulatory T cells (Treg) persist into adulthood where they play key roles in early protection from pathogens^12^ and suppression of organ-specific autoimmunity^13^, respectively. However, mechanisms regulating differentiation of functionally distinct T cell subsets in the perinatal versus adult thymus are not well understood.

Two major, non-mutually exclusive mechanisms could contribute to altered thymic function in the perinatal versus adult period. Age-associated changes in the hematopoietic compartment could drive altered T cell differentiation in the perinatal thymus. Fetal versus adult hematopoietic stem cells (HSCs) differ in their lineage potential^14,15^, with fetal HSCs biased towards production of T-lineage progenitors. Notably, the unique innate-like properties of perinatally derived CD8 T cells reflect their fetal HSC origin^8,12^. In addition, there is evidence that age-associated changes in the thymic microenvironment alter T cell differentiation. Genes involved in antigen processing and presentation are differentially expressed in perinatal versus adult thymic stromal cells^13,16^. Furthermore, the perinatal thymus produces elevated levels of type I and type III interferons (IFNs), which regulate antigen processing and presentation in thymic antigen presenting cells (APCs)^16,17^. Moreover, thymic epithelial cells (TECs) influence B cell differentiation in the perinatal thymus^18^. Because bi-directional crosstalk between thymic stromal cells and developing T cells has downstream consequences for T cell differentiation and selection, it is essential to determine how hematopoietic and stromal cell compartments change over the perinatal to juvenile transition.

Here we use single-cell transcriptional profiling to comprehensively evaluate the cellular composition and transcriptional profiles of thymic stromal cells from birth through the early juvenile period. These data reveal remarkably coordinated and dramatic changes in the cellular composition of TECs, mesenchymal cells, endothelial cells (ECs), and hAPCs coincident with the transition from perinatal thymic growth to juvenile homeostasis. Concurrent with these cellular changes, we identify transcriptional programs in each cellular compartment associated with age-associated growth and/or functional characteristics. We demonstrate that the transition in thymus growth is not regulated by the fetal to adult HSC switch, consistent with a stromal-intrinsic mechanism. The phenotype and function of thymocyte subsets, including Treg, change concomitantly with age-associated changes in the stromal compartment, indicating the essential role of the thymic microenvironment in regulating thymocyte development and selection. We identify parallel changes in growth characteristics of the human thymus and define the timing of the switch between growth and homeostasis. Together, these data define a perinatal to juvenile transition across all major thymic stromal and hAPC compartments and provide a resource to explore unique aspects of the perinatal versus juvenile thymus microenvironment.

## RESULTS

### A transition between perinatal thymus expansion and juvenile homeostasis occurs in mice and humans and is regulated by the cyclin D1-RB-E2F pathway in TECs

Thymus growth in mice increases exponentially after birth until postnatal day 10-14 (P10-P14) when growth plateaus, leading to a period of homeostasis (Figure 1A). The comparable pattern of thymus growth in males and females demonstrates that sex does not regulate this growth transition. The human perinatal thymus shows a comparable early expansion phase which transitions to homeostasis at ∼3-4 months of age in both sexes (Figure 1B). The mechanism(s) responsible for the shift from rapid thymus growth to homeostasis could be driven by the transition from fetal to adult hematopoiesis. To test this, we sustained Lin28 expression in Vav-iCre;R26iLin28a mice^19,20^, which results in a persistent fetal hematopoietic phenotype^19^. However, despite maintenance of Lin28 expression in hematopoietic cells through 9 weeks of age (Figure S1A), thymus size did not increase above control levels (Figure 1C). Thus, the switch from fetal to adult hematopoiesis does not drive the transition from thymus growth to homeostasis.

**Figure 1:**
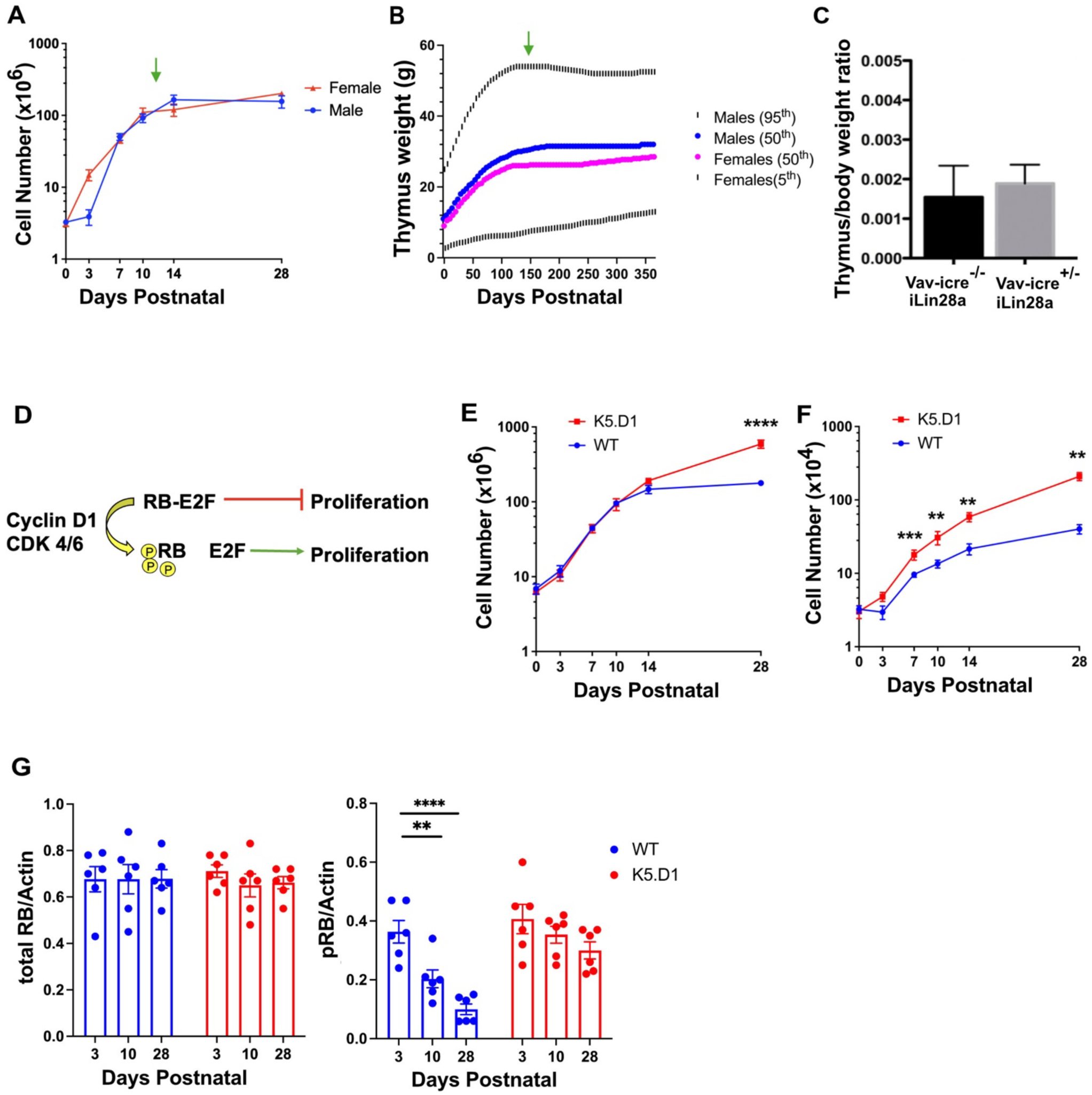
The Cyclin D1-RB-E2F pathway regulates the transition from perinatal thymus expansion to juvenile homeostasis. (A) Quantification of total thymus cellularity in male and female C57BL/6J mice at the indicated ages. Data are pooled from at least 3 experiments per age group (n=2-14 mice per sex per age group) and shown as mean ± SEM. Green arrow indicates the approximate age at which perinatal thymus growth transitions to homeostasis. (B) Quantification of human thymus weight at indicated ages post birth, as shown in previously published percentile charts^29^. Symbols represent thymus weight at 50th percentile for males (blue) and females (pink), black lines indicate thymus weight at 95th percentile (males) and 5^th^ percentile (females). n= 5633 males; n= 407 females. Green arrow indicates approximate age at which perinatal thymus growth transitions to homeostasis. (C) Thymus to body weight ratio in 8-week-old Vav-iCre^+/-^;R26iLin28a mice and Vav-iCre^-/-^;R26iLin28a control mice. Ratios are shown as mean ± SEM. (D) Simplified schematic of Cyclin D1-RB-E2F axis (E) Quantification of total thymus cellularity in WT and K5.D1 mice at indicated ages. Data are pooled from at least 3 experiments per age group (n=8-32 mice per genotype per age group) and shown as mean ± SEM. (F) Quantification of total TEC cellularity in WT and K5.D1 mice at the indicated ages. Data are pooled from at least 3 experiments per age group (n=8-32 mice per genotype per age group) and shown as mean ± SEM. (G) Quantification of western blot data showing total or phosphorylated Rb to actin ratios in WT or K5.D1 TECs. Each symbol represents data pooled from 6-8 individual mice and bars represent mean ±SEM from 6 independent experiments per age group per genotype. For Figures 1A, E and F, statistical analysis was performed using multiple t-tests; unpaired t-test per age group with FDR at 1% using two-stage step-up (Benjamini, Krieger, and Yekutieli). For Figure 1G statistical test was performed using one way ANOVA with Tukey’s multiple comparisons test. *P<0.05, **P<0.01, ***P<0.0001, and ****P< 0.00001.

We have shown that the thymus in RB deletion mutants and in mice expressing a keratin 5 promoter-driven cyclin D1 transgene (K5.D1) grows continually and fails to establish thymus homeostasis^21,22^. The K5.D1 transgene is expressed by TECs, but not by hematopoietic cells (Figure S1B), indicating that enforced expression of cyclin D1 in TECs is responsible for sustained thymus growth. Cyclin D1 activates cyclin dependent kinase 4 (Cdk4) and Cdk6, which phosphorylate and inactivate the retinoblastoma (RB) protein, releasing E2F transcription factors to promote cell cycle progression^23–28^ (Figure 1D). Thymus cellularity increases equivalently in wildtype (WT) and K5.D1 transgenic mice from birth until P10, after which the K5.D1 thymus continues to expand and fails to establish homeostasis (Figure 1E). Similarly, the number of WT TECs stabilizes after P14, whereas K5.D1 TEC cellularity continues to increase (Figure 1F). Notably, the divergence between WT and K5.D1 TEC cellularity occurs as early as P7, prior to the divergence in total thymus cellularity, reflecting the critical role of TECs in regulating thymus growth.

To determine if timing of the growth to homeostasis transition is linked to age-related regulation of RB activity in TECs, we assessed RB phosphorylation levels in P3, P10, and P28 WT and K5.D1 TECs (Figures 1G and S1C). Although total RB levels remained constant with age, the pool of phosphorylated RB decreased significantly after P10 in WT, but not in K5.D1 TECs. These results suggest that diminished RB phosphorylation over the transition rescues its ability to suppress proliferation and switch off growth in WT TECs to establish homeostasis, whereas maintaining RB phosphorylation in K5.D1 TECs blocks its inhibitory activity to prolong TEC proliferation and thymus growth.

### Single-cell RNA sequencing reveals dynamic changes in TEC transcriptional profiles and cellular composition across the transition

To determine if the perinatal to juvenile transition is accompanied by changes in TEC subset composition and/or transcriptional profiles, we performed single-cell RNA sequencing (scRNAseq) on sorted CD45^-^EpCAM^+^ WT TECs at P0, P3, P7, P10, P15 and P28 (Figure S2A). The concatenated scRNA-seq data were projected onto a Uniform Manifold Approximation and Projection (UMAP), and the major transcriptional clusters comprising cTECs, nurse cells, Ccl21^hi^ mTECs, Aire^hi^ mTECs and Aire^lo^ mTECs were annotated based on published gene expression signatures (Figures 2A and 2B)^30–38^. Nurse cells, which are cTECs that encompass thymocytes, have a mixed transcriptional signature of both cell types^39,40^. Clustering at higher resolution revealed additional transcriptional diversity (Figures S2B and S2C). For example, as previously reported, Aire^lo^ mTECs contain distinct subsets corresponding to mimetic TECs such as tuft cells, microfold cells, and ciliated cells^31–33,35,41,42^.

**Figure 2:**
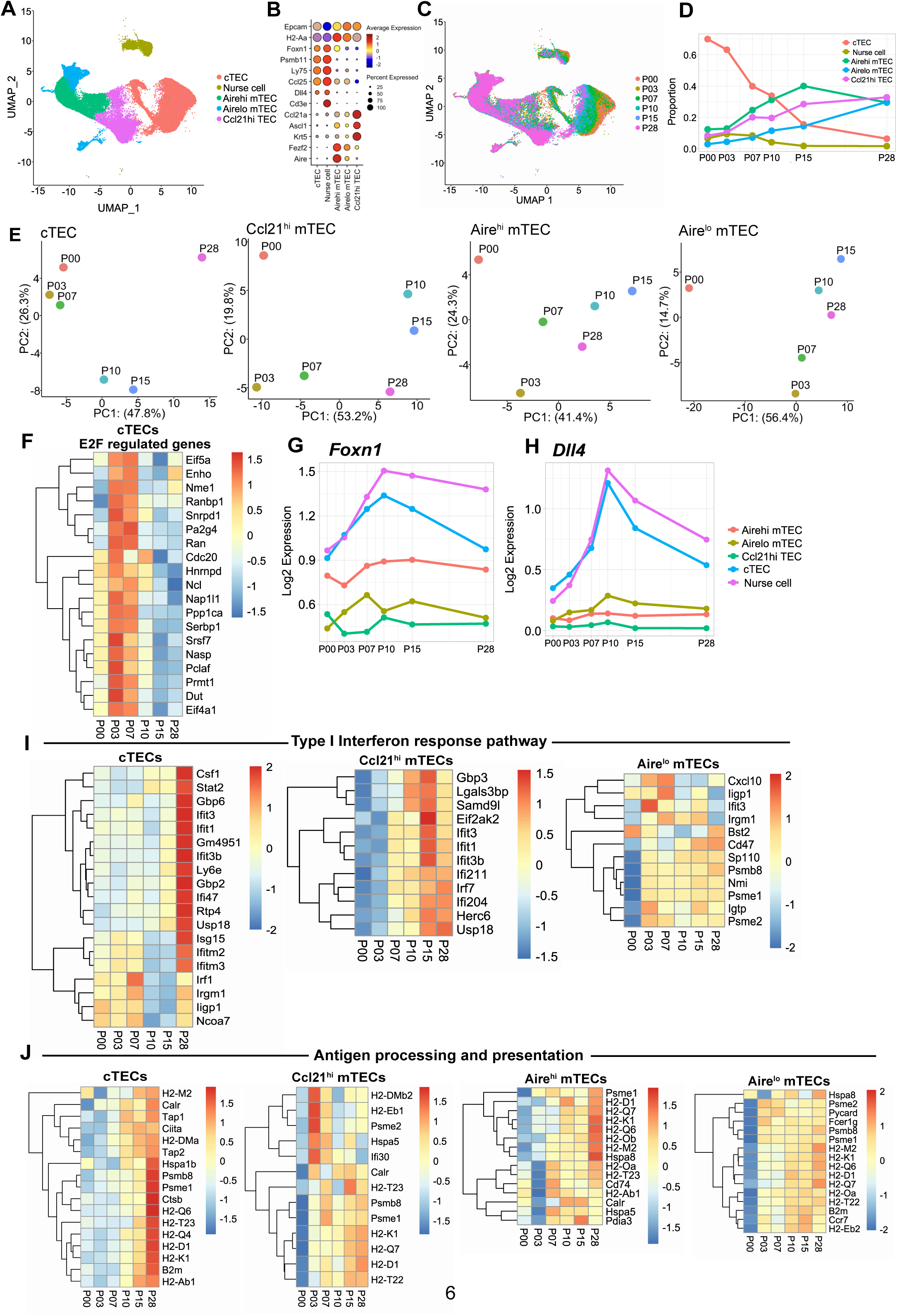
Changes in TEC transcriptional signatures and subset composition over the perinatal to juvenile transition. (A) UMAP visualizing scRNAseq data from 49,346 TECs from P0, P3, P7, P10, P15 and P28 C57BL6/J mice; n=2 for each timepoint except P15 (n=1). Colors indicate major TEC subsets. P denotes postnatal day. (B) Dot plot shows average expression of genes used to annotate TEC subsets in (A). (C) UMAP from (A) color-encoded by age. (D) Quantification of the frequency of TEC subsets within the total TEC compartment at the specified ages, based on scRNAseq datasets. (E) PCA plots show variance in gene expression across PC1 and PC2 in major TEC subsets. Colored symbols indicate age. (F) Heatmap showing normalized expression of E2F regulated genes, from GSEA of k-means clusters (see Figure S2G), in cTECs at the indicated ages. (G) Line graph showing normalized expression of *Foxn1* in major TEC subsets at the indicated ages. (H) Line graph showing normalized expression of *Dll4* in major TEC subsets at the indicated ages. (I) Heatmaps showing normalized expression of IFN responsive genes, from GSEA of k-means clusters (see Figure S2G), in TEC subsets. (J) Heatmaps showing normalized expression of antigen processing and presentation genes, from GSEA of k-means clusters (see Figure S2G), in TEC subsets.

The frequency and phenotype of cTECs and mTECs change markedly across the perinatal to juvenile transition. When scRNAseq data were color-coded by age a striking rainbow pattern emerged in cTECs, indicating they are particularly transcriptionally dynamic across the transition (Figure 2C). Principal component analysis (PCA) also showed that transcriptional variance in cTECs corresponded with age along the PC1 axis (Figure 2E). Moreover, P10 and P15 transitional age datasets were separated from earlier and later ages along the PC2 axis. Similarly, P10 and P15 Ccl21^hi^ mTEC datasets also clustered together on the PC2 axis (Figure 2E), together indicating distinct transcriptional states of TEC subsets at the transition. Both transcriptional and flow cytometric analyses showed that the relative abundance of cTECs declines while that of mTECs increases over the perinatal to juvenile transition (Figures 2D, S2D-S2F). The majority of both cTECs and mTECs have an MHCII^hi^ phenotype prior to P14, which switches to a predominantly MHCII^lo^ phenotype after the transition (Figure S2F).

We next performed k-means clustering of TEC datasets to group genes with similar age-associated expression patterns into modules (Figure S2G), and GSEA^43,44^ identified pathways enriched in each module. E2F-regulated target genes were enriched in cTEC and mTEC modules that peaked in neonates and declined over the transition (Figures 2F, S2G and S2H). For example, in cTECs, E2F-regulated gene expression peaked at P3-P7, then declined (Figure 2F). This decline in expression of E2F-regulated genes is consistent with reduced RB phosphorylation in TECs over the transition (Figure 1G), further implicating RB-mediated inhibition of E2F activity. Consistent with the presence of E2F binding sites in the *Foxn1* promoter^22^, expression of *Foxn1* and its target *Dll4* increased until P10 and declined thereafter (Figures 2G and 2H).

To identify additional factors that regulate age-associated transcriptional changes in TECs, we used the Encode database to perform transcription factor enrichment analysis on the top 200 variable genes specifying PC1 and PC2 (Figure 2E). In addition to *Foxn1*, transcription factors activated by IFN signaling (e.g. *Stat1* and *RelA*)^45,46^ were identified (Figure S2I). Recent reports have shown that IFNs are produced by mTECs and plasmacytoid dendritic cells (pDCs) and IFN signaling activates thymic APCs promoting central tolerance^16,17,47,48^. Consistent with the transcription factor enrichment, GSEA^43,44^ analysis revealed a progressive increase in expression of genes associated with the type I IFN response pathway in cTECs, Ccl21^hi^ mTECs and Aire^lo^ mTECs over the perinatal to juvenile transition (Figure 2I). IFN signaling promotes expression of genes involved in antigen processing and presentation^17^, and GSEA^43,44^ analysis also revealed an age-associated enrichment of antigen processing and presentation genes in the same TEC subsets in which the Type I IFN response pathway was activated (Figures 2J and S2G). Collectively these results suggest that altered antigen processing and presentation by TECs responding to IFN signaling may differentially regulate positive selection and central tolerance across the perinatal to juvenile transition.

To ascertain age-associated changes in thymus organization, we stained thymus sections with antibodies to keratin 5 (Krt5), Krt8 and Krt14. Distinct medullary islets containing Krt5+Krt14+ mTECs were present in P3-P7 sections (Figures S3A and S3B)^49^. Coalescence of these islets initiated during the P10-P14 transition and by P28 they had consolidated into central medullary regions as previously reported (Figure S3C)^50^. At P3 a high frequency of Krt8+Krt5+ cTECs were widely distributed between medullary islets but after P10 were primarily localized at the cortico-medullary junction (Figure S3B). The timing of these changes in organization of cTEC and mTEC subsets defined by keratin expression corresponded with the shift from rapid thymus growth to homeostasis.

### Single-cell RNA sequencing reveals changes in transcriptional profiles and composition of thymic mesenchymal, mesothelial and endothelial cells across the perinatal to juvenile transition

Mesenchymal and endothelial cells (ECs) in the thymus microenvironment play an essential role in the development and maintenance of TECs and thymocytes^51–53^. Therefore, we used scRNAseq to investigate changes in the cellular composition and transcriptional profiles of thymus mesenchymal and endothelial cells over the transition. Unsupervised clustering of CD45^-^ thymic stromal cells from WT mice yielded TECs, endothelium, mesenchyme, and mesothelium based on archetypal gene expression^54^ (Figures 3A and 3B). When color-encoded by age, the UMAP shows age-associated transcriptional changes in each stromal cell type (Figure 3C). Moreover, between birth and P14, mesenchymal cells decline in frequency, while TECs increase (Figure 3D). Notably, after peaking at P14, the proportion of endothelial cells in the stromal compartment declined by ∼50% by P28 (Figure 3D).

**Figure 3.**
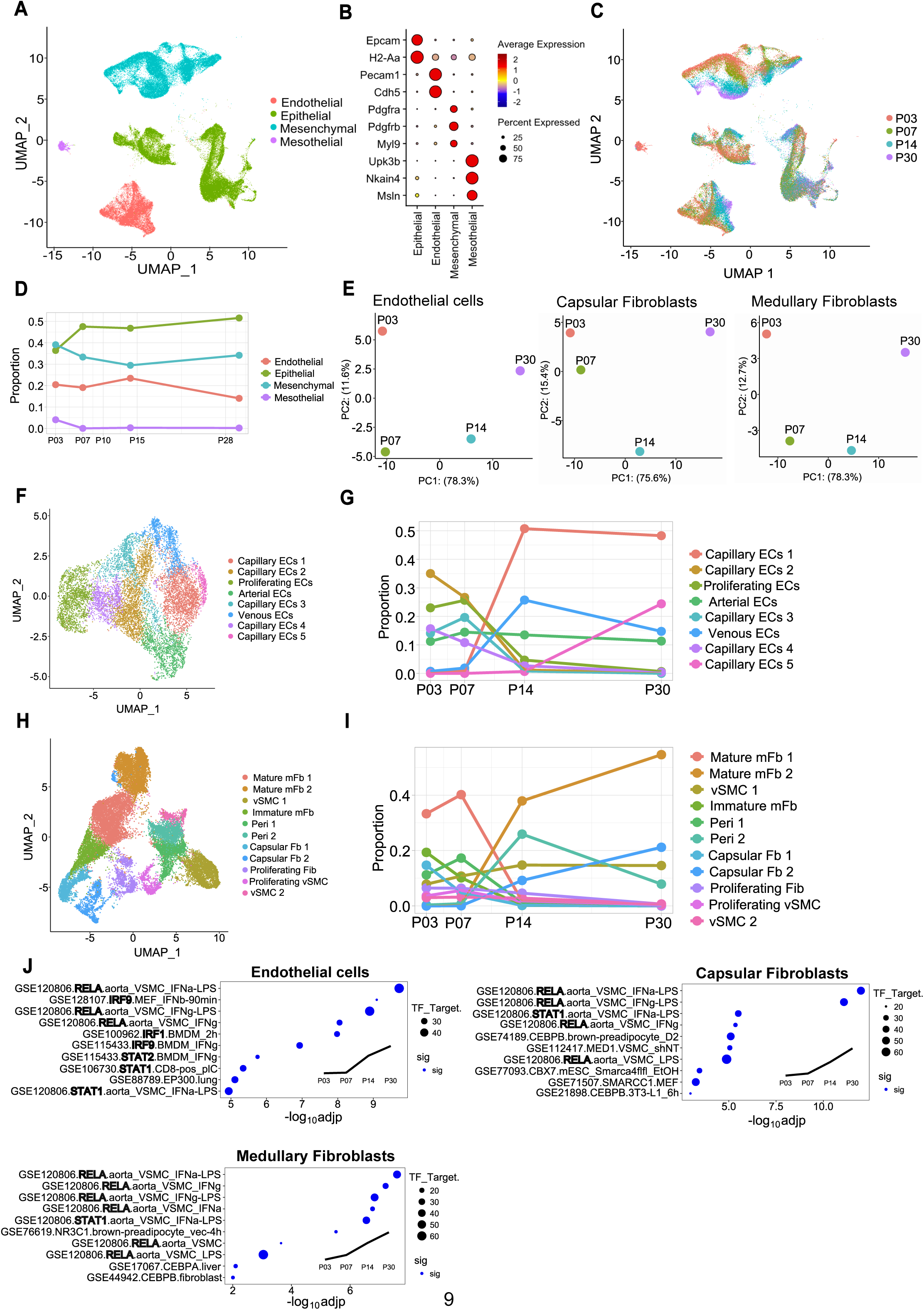
Endothelium and mesenchyme exhibit major transcriptome differences across the perinatal transition point. (A) Concatenated UMAP visualizing scRNAseq data from CD45^-^ total thymic stromal cells from P3, P7, P14, and P28 C57BL6/J mice, colored by major stromal subsets. (B) Dot plot showing expression of archetypal genes used to identify major stromal subsets. (C) UMAP visualization of the CD45^-^ total thymic stroma color-encoded by age. (D) Quantification of the proportions of major thymic stromal subsets within the total thymic stromal compartment at the indicated ages, based on scRNAseq datasets. (E) PCA plot visualization of pseudo-bulked endothelial cells, capsular fibroblasts, and medullary fibroblasts shows age-associated variation in gene expression in each subset. (F, H) UMAP plot visualization of subsampled scRNAseq data from (F) endothelium and (H) mesenchyme. (G, I) Quantification of the proportions of subsampled (G) endothelial and (I) mesenchymal subsets within the major cell types at the indicated ages. (J) The top 10 most significantly enriched transcription factors regulating a specific time-dependent gene expression pattern (shown in the subgraph) in endothelial cells, capsular fibroblasts and medullary fibroblasts. Node size represents the number of overlapping genes between the selected gene set and transcription factor binding targets. Nodes in blue indicate statistically significant enrichment (adjp < 0.05), while gray denotes non-significant enrichment.

To better understand the mesenchymal and endothelial data structure, we carried out PCA of the datasets from these stromal cell types. Higher resolution clustering allowed for separation of fibroblasts into capsular and medullary subsets based on established marker gene expression (Figure S4A)^51,55,56^. PCA revealed age was the main factor determining transcriptional variance along the PC1 axis, which separates datasets at P3 and P7 from those at P14 and P30 (Figure 3E). Furthermore, PC2 identified a distinct transcriptional profile of ECs, capsular fibroblasts, and medullary fibroblasts at the transitional ages of P7 and P14. Higher resolution clustering of endothelial cells (ECs) yielded arterial, venous, capillary, and proliferative ECs (Figures 3F and S4B). Higher resolution clustering of mesenchyme identified fibroblasts (Fb), vascular smooth muscle cells, capsular Fb, immature medullary fibroblasts (mFb)^51^, mature mFb, vascular smooth muscle cells (vSMC), and pericytes (Peri; Figure 3H). Notable age-associated changes occurred in the composition of ECs (Figure 3G) and mesenchymal cells (Figure 3I), with a progressive decrease in proliferative populations. Venous ECs became more abundant between P7 and P14. Capillary EC subsets 3, 4, and 2, which were abundant at P3 and P7, declined dramatically by P14, when subsets 1 and 5 become predominant (Figure 3G). The large separation of these capillary EC subsets on the UMAP indicates a major age-associated transcriptional shift. Similarly, in the mesenchymal compartment, immature medullary fibroblasts nearly disappear between P7 and P14 (Figure 3I). Capsular Fb, mature mFb, and Peri undergo large transcriptional shifts between P7 and P14, resulting in apparent compositional changes between subtypes of these subsets, such as mature mFB1 versus mature mFB2. We also carried out transcription factor enrichment analysis of the top 200 variable genes that defined PC1 and PC2 (Figure 3E). Strikingly, in ECs, cFbs, and mFbs, interferon responsive STAT1 and/or RELA were the major transcription factors associated with genes upregulated across the perinatal to adult transition (Figure 3J). In addition, as shown in Kang et al, 2025 (revision submitted), a severe decline in expression of *Igf2*, *Mest*, and *Dlk1* shaped the distinct transcriptomes of mesenchymal and ECs after P14. Together, these analyses identify IGF2 and IFN signaling as central transcriptional changes in the stromal compartment during the transition from perinatal thymus growth to juvenile homeostasis.

We previously reported that the frequency of subcapsular fibroblasts is severely reduced in the mouse thymus after the perinatal to juvenile transition (Kang et al, 2025, revision submitted). Similarly, the density of VIMENTIN+ human fibroblasts at the capsule was greatly reduced in the human thymus after the transition from perinatal expansion (3-day-old thymus) to juvenile homoeostasis (4-year-old thymus) (Figures S4C-S4F).

### scRNAseq reveals dynamic changes in cellular composition and molecular profiles of thymic hAPCs during the perinatal to adult transition in mice and humans

Because hAPCs participate in crosstalk with TECs, which undergo age-associated changes, and hAPCs induce thymocyte central tolerance to a wide variety of self-antigens^16,41,57–59^, we tested whether these key thymic APCs change in cellular composition and/or transcriptional profiles over the perinatal to juvenile transition. We carried out scRNAseq analysis of FACS purified hAPCs (Figure S5A) from WT mice at P0, P3, P7, P10, P14, and P28. A UMAP representation of the concatenated datasets reveals 4 major hAPC clusters, conventional dendritic cells (cDCs), plasmacytoid DCs (pDCs), B cells, and macrophages, which were annotated based on expression of established lineage specifying genes (Figures 4A and 4B). Different hAPC subsets induce tolerance to distinct categories of self-antigens. cDCs induce tolerance to self-antigens from their own proteomes or acquired from peripheral tissues, circulation, or mTECs^58,60–64^. B cells undergo CD40-dependent licensing, likely enforcing tolerance to self-antigens expressed by activated B cells in the periphery^58,59^. The contributions of macrophages and pDCs to central tolerance remain less well-defined, though pDCs have been implicated in inducing tolerance^65^. The cellular composition of hAPCs becomes altered over the perinatal to juvenile transition (Figures 4C, S5B and S5C). The frequencies of B cells and pDCs increase after birth (P0), peaking at P10-P14 transitional ages. cDCs decline in frequency from birth through transitional ages but rebound by P21-P28. Macrophage frequencies decline between P0 and transitional ages (P10-P14). Notably, B cells are the predominant hAPC subset during the transition period. In human thymus, the frequency of B cells within the hAPC compartment also increases between the perinatal (≤2 months) and post-transition (7-14 month) ages, and continues to increase through at least 4-8 years (Figures S5D and S5E).

**Figure 4:**
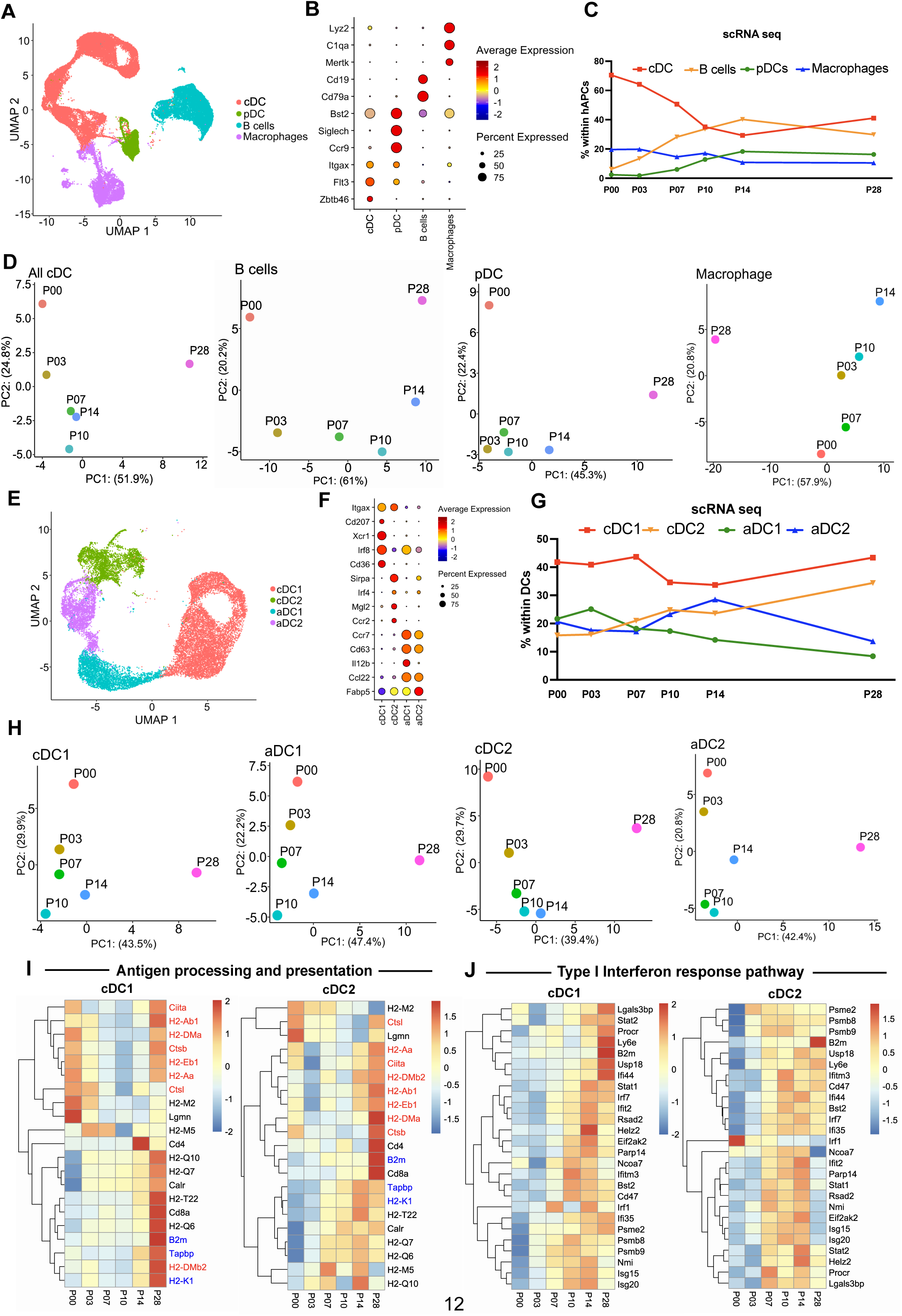
scRNAseq reveals changes in hAPC and DC composition and transcriptional profiles over the transition. (A) Concatenated UMAP visualizing scRNAseq data from 40,913 thymic hAPCs from P0, P3, P7, P10, P14, and P28 C57BL6/J mice (n=2 per age). Colors indicate annotated cell subsets. (B) Dot plot shows average normalized expression values of genes used to annotate hAPC subsets. (C) Quantification of cDCs, B cells, pDCs and macrophages as a frequency of total hAPCs at the specified ages, based on scRNAseq data. (D) PCA plots showing variance in gene expression across PC1 and PC2 in the indicated hAPC subsets. Colored symbols indicate age. (E) UMAP visualization of higher resolution clustering of scRNAseq datasets from the 20,401 thymic cDCs, from Figure 4A. (F) Dot plot shows average normalized expression values of genes used to annotate cDC subsets. (G) Quantification of cDC1s, cDC2s, aDC1s, and aDC2s as a frequency of total DCs at the specified ages, based on scRNAseq data. (H) PCA plots showing variance in gene expression across PC1 and PC2 in the indicated cDC subsets. Colored symbols indicate age. (I) Heatmaps showing normalized expression of genes associated with the antigen processing and presentation pathway, from GSEA of k-means clusters (see Figure S6F), in cDC1 and cDC2 subsets at the indicated ages across the perinatal to juvenile transition. Genes highlighted indicate MHCI genes (blue) and MHCII genes (red). (J) Heatmaps showing normalized expression of genes associated with the Type I IFN response pathway, from GSEA of k-means clusters (see Figure S6F), in cDC1 and cDC2 subsets at the indicated ages across the perinatal to juvenile transition.

We carried out PCA to determine if age was a key contributor to gene expression variability in the four major hAPC subsets. PC1, which accounts for 45-50% of the variation in gene expression, corresponded with age in cDCs, B cells, and pDCs (Figure 4D). Moreover, PC2, which captures >20% of gene expression variation in B cells, pDCs and cDCs separates P0 and P28 ages from the transitional ages (P7, P10, P14), suggesting that thymic hAPCs undergo a distinct transcriptional shift during the transition from perinatal growth to juvenile homeostasis.

### scRNAseq reveals dynamic changes in transcriptional profiles of thymic DCs over the perinatal to juvenile transition

To elucidate the biology underpinning age-associated changes in cDCs, scRNAseq data were first clustered at higher resolution, revealing four major subsets, cDC1s, cDC2s and activated cDC1s and cDC2s, denoted ‘aDC1s’ and ‘aDC2s’, respectively (Figure 4E), with annotations based on lineage-specifying genes (Figure 4F)^66,67^. In agreement with prior reports^66,68,69^, expression of *Itgax* declines upon thymic DC activation, as does expression of *Xcr1* and *Sirpa* by aDC1s and aDC2s, respectively. Additional DC heterogeneity was revealed by clustering cDCs at higher resolution (Figures S6A and S6B). When scRNAseq data were color-encoded by age, transcriptional shifts across the transition were most evident in cDC1s (Figure S6C). We tested if cDC subset composition changes over the perinatal to adult transition by analyzing scRNAseq (Figure 4G) and flow cytometry data (Figures S6D and S6E). Intrathymically derived cDC1s are the predominant DC subset in the perinatal P0 mouse thymus. cDC1 frequencies decline slightly at transitional ages (P10-14) when cDC2s, which migrate into the thymus^60,70–73^, and downstream aDC2s become more frequent. Based on scRNAseq, aDC1 and aDC2 are present at P0, and aDC1 frequencies decline steadily over the perinatal to juvenile transition (Figure 4G). However, aDCs were not detected by flow cytometry at P0 (Figure S6E), which reflects the lack of cell surface expression of CCR7 and CD63, phenotypic markers that delineate aDCs (Figure S6D), on perinatal cDCs, despite expression of *Ccr7* and *Cd63* transcripts. These findings demonstrate the importance of evaluating subset composition through phenotypic and transcriptional approaches to obtain a comprehensive assessment of age-associated changes. Given that aDC1s efficiently acquire and present mTEC-derived antigens to thymocytes^62,64,67,69^, while cDC2s efficiently induce Treg generation in vitro^72,74^, the age-associated changes in frequencies and transcriptional profiles of DCs could alter central tolerance, contributing to selection of functionally distinct T cells across the perinatal to juvenile transition^7,9,13^.

To evaluate age-associated transcriptional changes in cDCs (Figures 4D and S6C), PCA analysis was performed on the four cDC subsets (Figure 4H). P28 data separated from perinatal and transitional datasets along the PC1 axis, which accounts for >39% of the transcriptional variance between samples in each subset. Notably, the PC2 axis separated transitional age DCs (P7, P10 and P14) from those at P0 and P28. Thus, adult DC subsets are transcriptionally distinct from perinatal DCs, and a unique transcriptional profile emerges in DCs at transitional ages between perinatal growth and homeostasis. We performed k-means clustering to identify modules of genes whose expression changes coordinately with age (Figure S6F), and GSEA^43,44^ revealed biologic pathways associated with each module. Notably, genes from the Antigen Processing and Presentation pathway as well as the Type I IFN Response pathway were enriched in multiple k-means clusters (Figure S6F). Recent studies show that Type I and III IFN signaling, which are both represented by the Type I IFN pathway signature^75,76^, are required for DC activation, Treg selection, and tolerance to self-antigens expressed under inflammatory conditions^16^. In both cDC1s and cDC2s, genes involved in MHCI antigen processing and presentation increase in expression starting at transitional ages and peaking at P28; in contrast, MHCII antigen processing and presentation genes declined in expression at transitional ages relative to P0 and P28 (Figure 4I). These findings indicate that age-associated changes in antigen processing and presentation on MHC-I versus MHC-II are temporally distinct in cDCs, with largely similar patterns maintained in activated DCs (Figure S6G). Many genes associated with the Type I IFN pathway, including proteasome components and *B2m*, which are key players in MHCI antigen processing and presentation, increase in expression in all DC subsets starting at transitional ages (P7-P14) and are maintained at P28 (Figures 4J and S6H), mirroring expression of MHCI antigen processing and presentation genes (Figure 4I).

To identify transcriptional regulators that could account for the age-associated transcriptional signatures, we performed transcription factor binding enrichment analysis of the top 200 variable genes contributing to PC1 and PC2 using the ENCODE database (Figure S6I). This analysis revealed that transcription factors activated by IFN signaling, such as STAT1, STAT2, IRF1, IRF8 and IRF9 were upregulated with age particularly at the transitional ages in cDC1 and cDC2. Altogether, these data indicate that IFN signaling is initiated in DCs at transitional ages, likely resulting in altered presentation of self-antigens in cDC1s and cDC2s and impacting thymocyte central tolerance.

### scRNAseq reveals changes in B cell composition and transcriptional signatures over the perinatal to adult transition

Thymic B cells are increasingly recognized as essential APCs that induce central tolerance^59,75,77^. Additional B cell subsets were identified through higher resolution clustering of B cell scRNAseq data, revealing 5 major subsets (Figure 5A) Cycling B cells were defined by expression of *Mki67* and *Top2a* (Figure 5B). B cell progenitors expressed *Rag1*, *Ebf1*, and *Vpreb3*, consistent with B cell differentiation in the thymus^77,78^. Immature B cells were identified as *Ighm^+^ Ighd*^lo^ cells with high levels of B220 (*Ms4a1*), while mature naïve B cells were *Ighm^+^ Ighd^hi^*. Activated B cells included class-switched B cells (*Ighm*^-^ *Ighd*^-^ cells expressing *Ighg2b* and *Igha)* and non-class-switched B cells (*Ighm^+^ Ighd^-^*) and expressed high levels of MHCI and MHCII genes and IFN-stimulated genes like *Ly6a and Ifi27l2a*. When color-encoded by age, the UMAP revealed an age-associated transcriptional shift in B cell subsets, with progenitors, immature B cells, and cycling B cells predominating at P0, followed by a transition first to mature naïve and then activated B cell subsets (Figures 5C and 5D).

**Figure 5:**
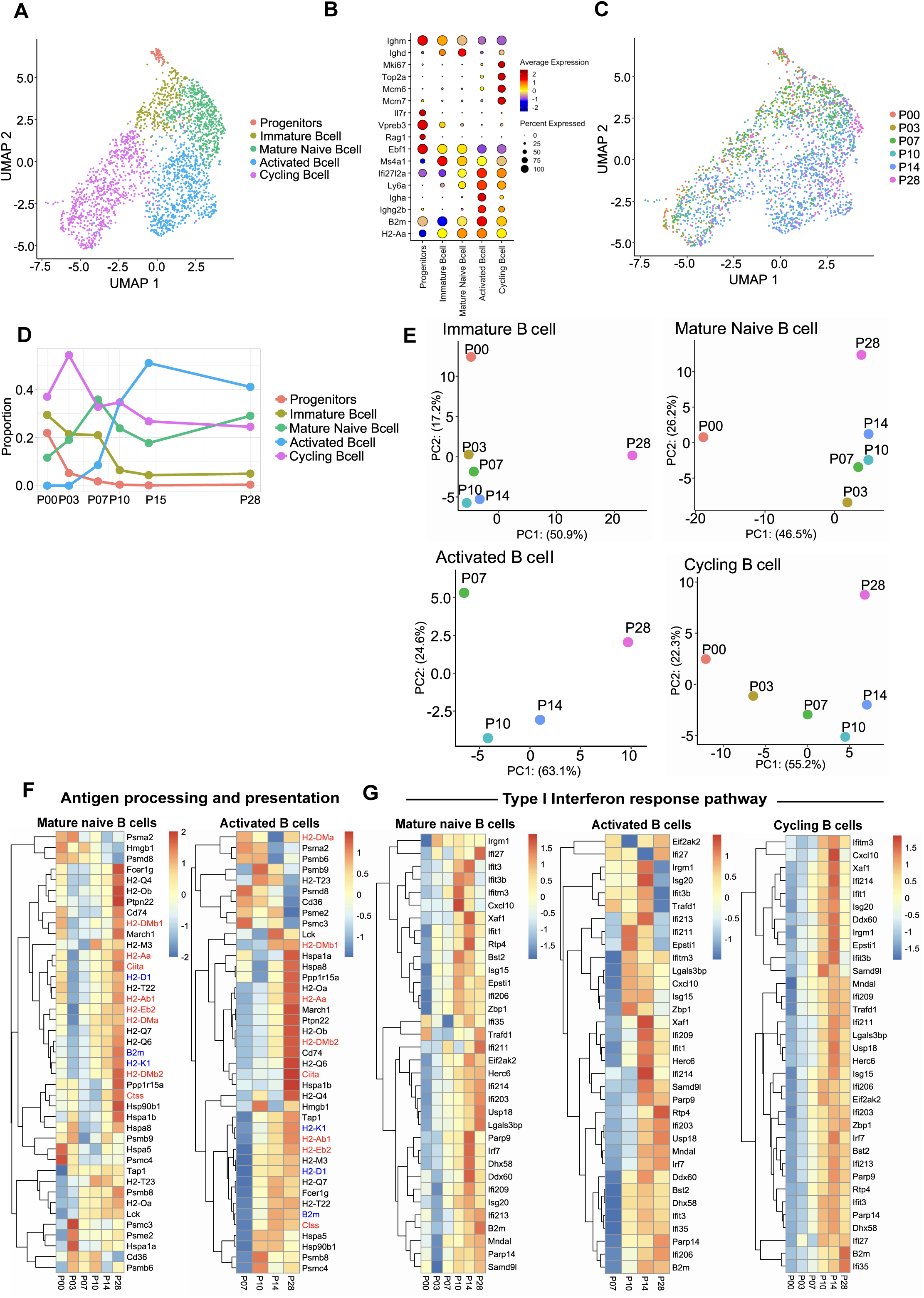
scRNAseq analysis shows changes in thymic B cell composition and transcriptional signatures over the perinatal transition. (A) Concatenated UMAP representation of scRNAseq data from thymic B cell subsets from P0, P3, P7, P10, P14, and P28 C57BL/6J mice (n=2 per age). (B) Dot plot shows normalized expression values of representative genes used to annotate B cell clusters. (C) UMAP from (A) color-encoded by age. (D) Quantification of the indicated B cell subsets as a frequency of total B cells at the specified ages, based on scRNAseq data. (E) PCA plots showing variance in gene expression across PC1 and PC2 in the indicated B cell clusters. Colored symbols indicate age. (F) Heatmaps showing normalized expression values of genes associated with the antigen processing and presentation pathway, from GSEA of k-means clusters (see Figure S7A), within mature naive and activated B cell subsets at the indicated ages. Genes highlighted indicate MHCI genes (blue) and MHCII genes (red). (G) Heatmaps showing normalized expression values of genes associated with the Type I Interferon response pathway, from GSEA of k-means clusters (see Figure S7A), within mature naïve, activated B cell, and cycling B cell subsets at the indicated ages.

PCA analyses revealed that PC1 stratifies cycling and activated B cell subsets by age (Figure 5E). Notably, PC2 corresponds with a unique transcriptional signature of the transitional period for B cell subsets (Figure 5E). Similar to age-associated transcriptional changes in TECs and DCs, genes associated with the Antigen Processing and Presentation as well as the Type I IFN response pathways were enriched in multiple k-means clusters of temporally coordinated gene expression (Figure S7A). Genes associated with MHCI processing and presentation, increased in expression in B cell subsets starting at transitional ages, while genes associated with MHCII antigen processing and presentation declined at P3 before increasing at transitional ages through P28 in mature naïve and activated B cell subsets (Figures 5F and S7B). Moreover, Type I IFN response genes also increased at transitional ages in mature naive, activated, and cycling B cell subsets (Figures 5G and S7C). Consistent with these findings, transcription factor enrichment analysis of the top 200 most variable genes in PC1 and PC2 shows that genes regulated by IFN-activated transcription factors STAT1, STAT2, IRF1, IRF8, and IRF9 increase in B cells during the transition, peaking in the activated B cell subset at transitional ages (Figure S7D). Expression of the Type I IFN-regulated transcript *Ly6a* increased with B cell maturation (Figure 5B), and type III IFN signaling has been shown to be required to activate thymic B cells, which results in Treg selection^75^. These findings indicate that IFN signaling is initiated in perinatal B cells and peaks at transitional ages in the activated subset, likely altering antigen presentation on MHCI and MHCII over the perinatal to juvenile transition, with consequences for age-associated changes in central tolerance.

### Thymocytes experience altered TCR signaling during the perinatal to adult transition, with intact clonal deletion, but altered selection of functionally distinct Tregs

Because our data reveal profound changes in the frequencies and expression profiles of all major APC subsets over the perinatal to juvenile transition, we evaluated whether temporally coordinated changes occur in thymocyte differentiation (Figure 6A). While cellularity of all thymocyte subsets increased with thymus growth, the frequency of DN (CD4^-^CD8^-^) thymocytes was highest at P0 and declined sharply between P3 and P7, as post-positively selected CD3^lo^CD69^+^ double positive (DP) and single positive (SP) thymocytes increased in frequency (Figures 6B and S8A). The timing of increased frequencies of post-positive selection thymocyte subsets correlates with the increased frequency of mTECs relative to cTECs (Figure S2E).

**Figure 6:**
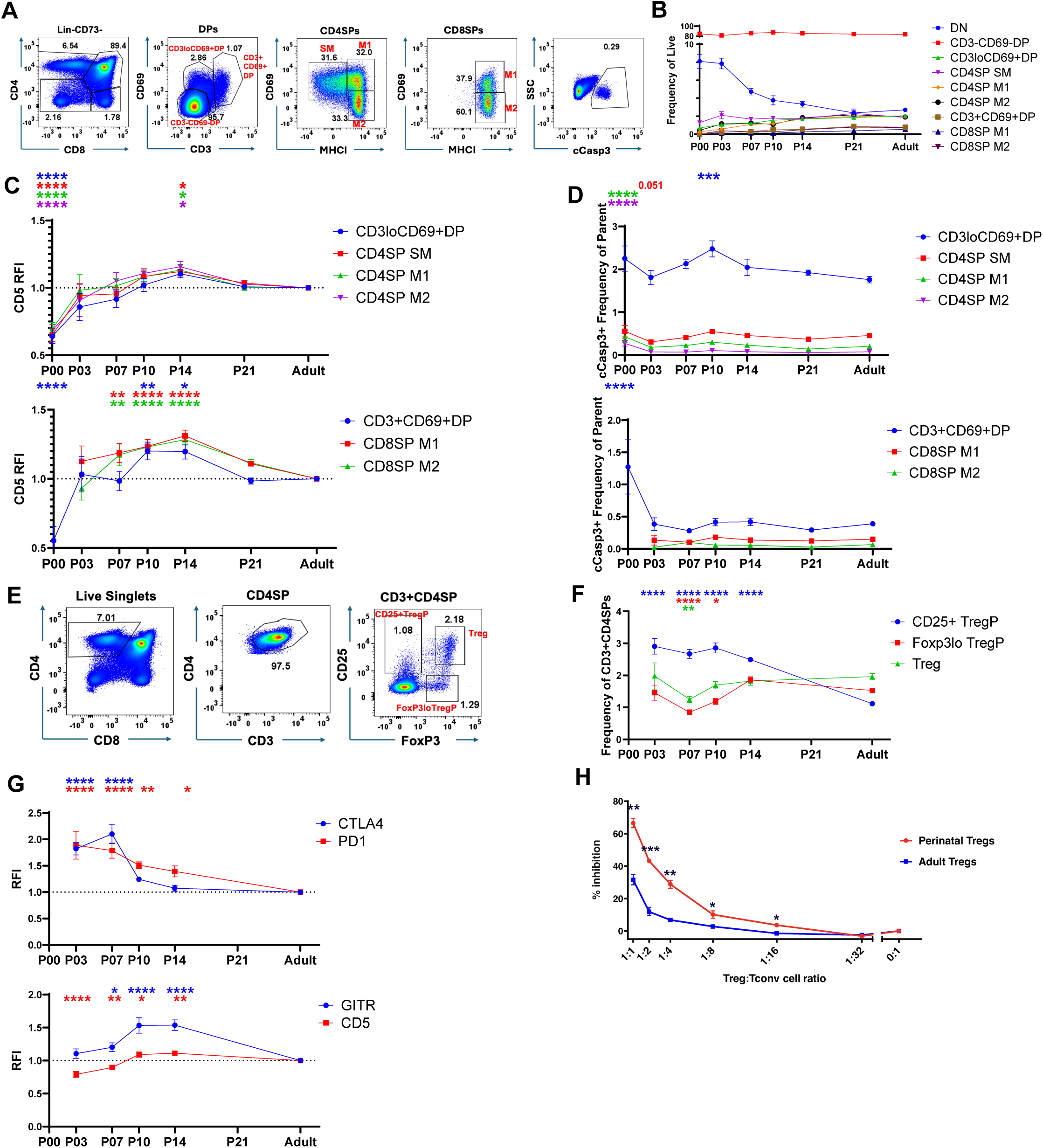
Conventional Thymocyte and Treg composition, phenotype, and function are altered at distinct stages over the perinatal to juvenile transition. (A) Representative flow cytometry data of thymocytes, delineating progressive stages of T cell development. The final plot shows representative gating on cleaved Caspase 3+ (cCasp3+) cells to quantify negative selection. Cells are pre-gated on live, singlet, lineage- (lin-), CD73-cells. (B) Quantification of thymocyte subsets as a frequency of live cells at the indicated ages. Data are pooled from at least 3 independent experiments per age group (n=8-47 mice per age group). Symbols with error bars represent mean ±SEM. (C) Quantification of CD5 expression levels on post-positive selection DP and CD4SP thymocytes (top) and CD8SP thymocytes (bottom) of the indicated ages (n=2-5 mice per age) relative to the average expression of at least 2 1-2mo adults within each experiment. Symbols with error bars represent mean ±SEM (D) Quantification of the frequency of cleaved cCasp3+ cells in each thymocyte subset at the indicated ages. Data are pooled from at least 3 independent experiments per age group (n=8-47 mice per age group). Symbols with error bars represent mean ±SEM. (E) Representative flow cytometry gating used to identify CD25+TregP, Foxp3loTregP, and CD25+Foxp3+ (Treg) within the CD4SP compartment. (F) Quantification of the frequency of Treg and TregP subsets with the CD4SP compartment, as shown in (E). Data are pooled from at least 3 independent experiments per age group (n=9-30 mice per age group). Symbols with error bars represent mean ±SEM (G) Quantification of expression levels of proteins associated with Treg suppressive capacity (CTLA4 and PD1 (top), GITR and CD5 (bottom) are shown relative to the average of at least 2 1-2mo adults within each experiment. Data are pooled from at least 3 independent experiments per age group (n=9-30 mice per age group). Symbols with error bars represent mean ±SEM (H) Percent (%) of Tconv cells inhibited from proliferating in the presence of P10 or adult splenic Tregs at the indicated Treg:Tconv cell ratios. Data are representative of n=4 independent experiments. Bars represent mean ± SEM and symbols represent average of triplicate wells at each Treg:Tconv cell ratio. For figures 7B-7F, an ANOVA with Dunnett’s multiple comparison correction to adults within each cell type was performed. For figure 7G, multiple unpaired t-tests with the Holm-Šídák method was performed. For 7H, statistical analysis was performed using Student *t*-test; *p<0.05, ** p<0.01, ***p<0.001, ****P<0.0001.

We next evaluated whether post-positive selection thymocytes differ qualitatively during the perinatal to adult transition. It was previously reported that CD4SP and CD8SP subsets express elevated levels of CD5 at transitional ages^79–81^, indicating an increased affinity of positively selected TCRs for self-antigens. We also found elevated CD5 expression at transitional ages P10 and P14 relative to P28; however, CD5 levels are quite low at P0 and are not elevated above P28 levels at P3 or P7 (Figure 6C). Furthermore, elevated CD5 expression occurs coordinately in all post positive-selection subsets, from the least mature CD3^lo^CD69^+^ DP cells to the most mature CD4SP M2 cells. If positive selection at the DP stage resulted in elevated CD5 levels, increased CD5 expression by SP thymocytes should lag temporally as CD5^hi^ DP thymocytes differentiate into CD4SP and CD8SP stages over a period of at least 4-5 days^47,82,83^. Thus, the coordinate increase in CD5 levels in all post-positive selection thymocyte subsets at P10 and P14 suggests the increase in CD5 may reflect a change in the thymus environment at transitional ages that increases the strength of TCR signaling in all thymocytes simultaneously. Because elevated CD5 levels indicate increased self-reactivity, we tested if the frequency of thymocytes undergoing negative selection declines at transitional ages but did not find evidence for this based on the frequency of cleaved caspase 3 TCR-signaled thymocytes. Instead, the frequency of post-positive selection DP thymocytes undergoing negative selection increases significantly at P10, with a similar trend in other subsets (Figure 6D). Thus, reduced negative selection does not account for the increased CD5 levels in post-positive selection thymocytes at transitional ages, in agreement with a prior study^79^.

In contrast to Treg selected in the adult thymus, those selected in the perinatal thymus display an elevated activation phenotype, persist into adulthood, and are able to suppress autoimmunity driven by *Aire*-deficiency^13,84^. Furthermore, expression of *Aire* uniquely in the perinatal period is necessary and sufficient to prevent autoimmunity suggesting that *Aire* expression by perinatal mTECs is required to select functionally unique Treg that protect against autoimmunity throughout life. However, it is unclear whether Treg selected in the perinatal thymus are endowed with more suppressive capacity than those selected in the adult thymus, or whether only perinatal Treg with greater suppressive activity persist into adulthood. Thus, we investigated Treg differentiation and function in the perinatal versus juvenile thymus. The frequency of CD4SP Treg remains relatively constant from P3 through P28, although numbers increase with thymus growth (Figures 6E, 6F and S8B). However, the frequency of CD25^+^Treg precursors (TregP), which are generated when CD4SP cells experience a strong TCR signal, is highest from P3-P14, consistent with a microenvironment that induces strong TCR signals at transitional ages (Figure 6F). Treg selected at P3 and P7 expressed the highest levels of PD1 and CTLA4, both of which are associated with heightened suppressive capacity^85,86^ (Figure 6G). In contrast, GITR, which is highly expressed on more autoreactive Treg precursors^87^, and CD5 are expressed at the highest levels at P10 and P14 (Figure 6G), when CD5 levels are generally highest on all post-positive selection thymocyte subsets (Figure 6C), indicating that Treg with distinct phenotypes are selected in the early perinatal versus transitional ages, both of which differ from juveniles (P28). We also found that Treg at a transitional age suppressed polyclonal T cell proliferation in vitro more efficiently than adult Treg (Figure 6H). Together, these data suggest that changes in thymic APCs at P7-14 result in selection of thymocytes and Treg with unique phenotypes and functional characteristics.

### Age-associated Expression of IFN responsive genes and Myc and E2F target genes is globally coordinated in diverse cell types in the perinatal thymus microenvironment

To identify common biologic processes altering gene expression in multiple cell types over the perinatal to juvenile transition, we performed tensor decomposition analysis on combined scRNAseq datasets from TECs and hAPCs (Figure 7A). The top Hallmark gene sets showing shared temporal expression patterns across TEC and hAPC subsets were IFN alpha response, MYC target genes and E2F target genes (Figure 7B). Heatmaps showing expression levels of the top 20 genes contributing to the shared temporal expression patterns showed that IFN alpha responsive genes increase, whereas MYC and E2F target genes decrease as the thymus transitions from perinatal growth to juvenile homeostasis (Figures 7C-7E). Although scRNAseq data from mesenchymal cells and endothelial cells were not included in the tensor analysis, because they were sampled at fewer ages, heatmaps showed the same temporal expression patterns of IFN alpha responsive and E2F target genes in these cell types (Figures 7F and 7G). Collectively, these data show that highly diverse cell types in the perinatal thymic microenvironment undergo remarkably concordant age-associated changes in gene expression reflecting diminished growth and increasing inflammatory stimuli.

**Figure 7:**
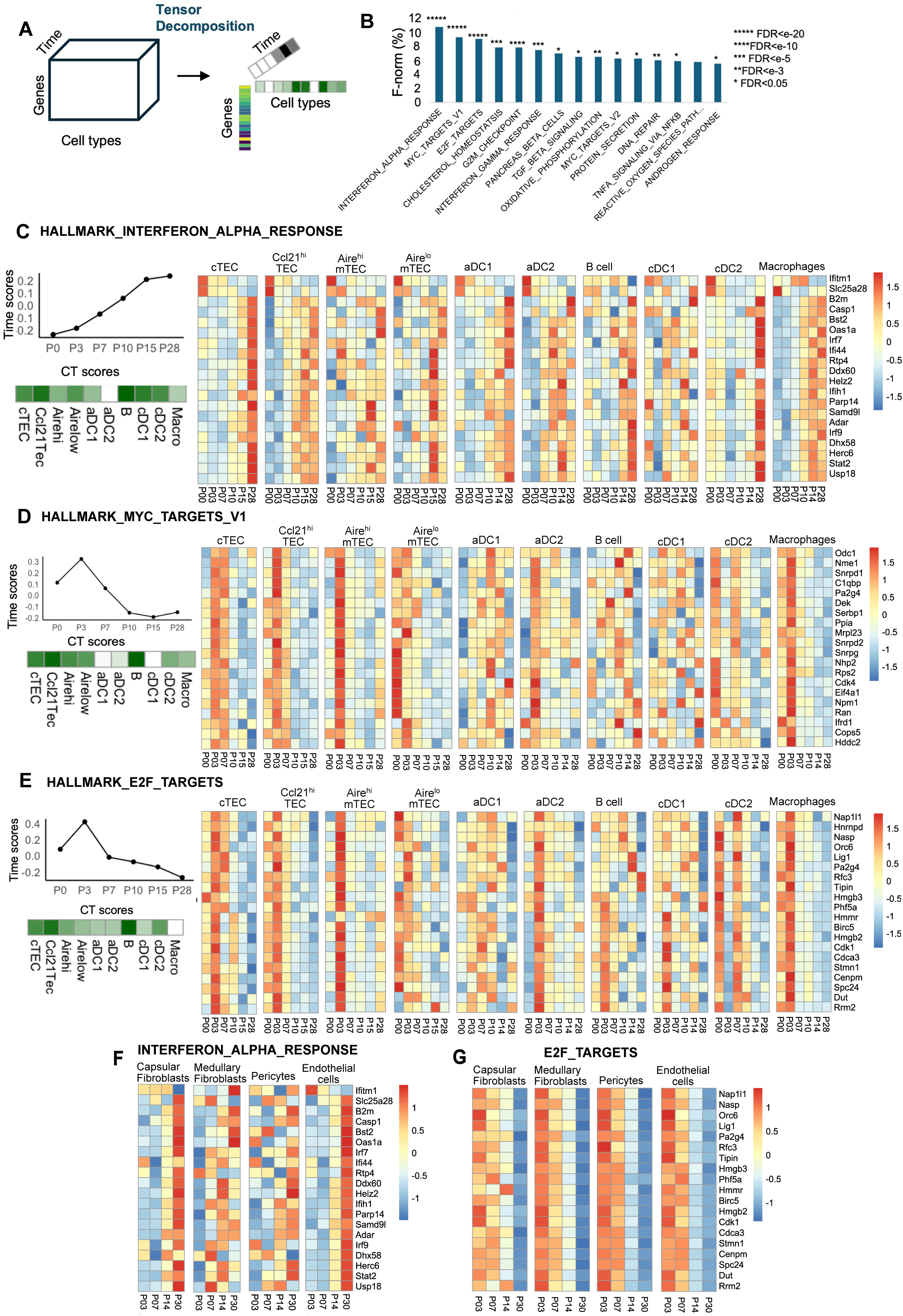
Type I IFN signaling, Myc targets, and E2F targets exhibit shared temporal expression patterns in highly diverse thymic cell types across the transition. (A) Schematic representation of tensor decomposition of hAPC and TEC scRNAseq datasets. (B) The top 15 Hallmark gene sets with pronounced temporal expression patterns across thymic hAPC and TEC subsets, with the percentage of the Frobenius norm (F-norm) indicating the total variance captured by each pattern. (C-E) For the Hallmark gene sets Hallmark_interferon_alpha_response (C), Hallmark_MYC_targets_v1 (D), and Hallmark_E2F_targets (E), the shared temporal expression patterns (Time scores) are presented, alongside cell type scores (CT scores), reflecting the strength of the pattern in each cell type, and heatmaps displaying the actual expression patterns of the 20 most contributing genes for thymic TEC and DC datasets at the indicated ages. (F) Heatmaps show the expression patterns of the 20 most contributing genes to the Hallmark_interferon_alpha_response gene set in thymic stromal cell subsets of the indicated ages. (G) Heatmaps show the expression patterns of the 20 most contributing genes to the Hallmark_E2F_targets gene set in thymic stromal cell subsets of the indicated ages.

## DISCUSSION

We have generated high-resolution, coordinated scRNAseq datasets with which to investigate changes in the transcriptional landscape and subset composition of TECs, mesenchymal cells, ECs and hAPCs in the thymus microenvironment as it shifts from perinatal growth to perinatal homeostasis. The results reveal a precisely timed process in which transcriptomic changes in diverse cell types parallel the shift from exponential thymus growth in newborns to homeostasis in juveniles. Notably, these transcriptomic changes are not merely a steady continuum of age-associated transitions, but reveal unexpected unique transcriptomic states for mesenchymal, hAPC, and TEC subsets that differ from either newborn or juvenile states. Moreover, we show that transcriptional and compositional changes observed in thymic APCs correspond temporally to phenotypic and functional changes in perinatal thymocytes. Together, these data sets provide a resource for further investigations into mechanisms that regulate age-associated shifts in growth and gene expression profiles of cells that comprise the perinatal thymic microenvironment and will facilitate understanding of how such changes influence development and selection of perinatal T cells including highly suppressive T cells which persist into adulthood^13^.

We previously reported that either deleting RB or blocking its activity in K5.D1 TECs prevents age-associated thymus involution^21,22^. Here we show that the Cyclin D1-RB-E2F pathway is also a key mechanism responsible for the precisely timed arrest in perinatal thymus growth. Because the WT and K5.D1 thymus have identical growth rates for approximately the first 10 days after birth, the initial rapid expansion in thymus size is likely independent of RB-mediated control. However, between P10 and P14 growth plateaus in the WT, but not in the K5.D1 thymus, suggesting that RB normally exerts its growth inhibitory activity during the perinatal transitional window. This inference is supported by a decline in the pool of phosphorylated RB in WT perinatal TECs, which does not occur in the continually expanding K5.D1 TECs and is consistent with an age-associated decrease in expression of E2F-regulated genes in WT TECs.

There is abundant evidence that TECs control much of the dynamics of thymic growth, function, and involution across the lifespan^50,88,89^. The key role of the RB pathway in regulation of *Foxn1*^22^ and the changes shown here in TEC subsets during the perinatal to juvenile transition indicate that TECs also control this milestone in the thymic lifecycle. Nonetheless, the shifts that also occur in the transcriptomes of endothelial and mesenchymal populations are at least as dramatic as those in TECs and indicate that functional changes in these populations are likely significant components of the shift to homeostasis. As there is also abundant evidence that TECs control the development, organization, and function of other stromal cell types in both the fetal and postnatal thymus, it is also likely that crosstalk drives these changes in non-TEC stroma. Notably, we have shown that both the structure and function of the fetal thymic vasculature and the interaction between endothelial and mesenchymal cells is directed by TECs^90^. Signals from non-TEC stroma can also feedback to TECs. cTEC regeneration after damage is influenced by BMP4 signaling from endothelial cells to cTECs^91^. Our data here strongly indicate that a decline in IGF2 signaling from capsular fibroblasts and endothelial cells is the major transcriptional change in these populations at the transition. Since IGF2 is known to regulate the RB-E2F pathway via CyclinD1 transcription^92,93^, it is a reasonable hypothesis that IGF2 from these populations acts on TECs to regulate the RB pathway upstream of Foxn1. The precipitous loss of IGF2 signaling at the transition could thus be the trigger that induces the change in RB activity and Foxn1 expression. However, the initiating event that triggers this drop in IGF2 expression across multiple cell types is unclear and could initiate from TECs themselves.

Across the perinatal to juvenile transition, hAPCs undergo major compositional changes that likely impact central tolerance. We observe a decline in the frequencies of cDCs and macrophages from birth through the transitional period, accompanied by an increase in the frequencies of both B cells and pDCs. Interestingly, B cells are the most abundant hAPC subset at transitional ages, suggesting that they may play a unique role in thymocyte selection in perinates. Thymic B cells undergo CD40-mediated activation/licensing through interactions with CD4SP thymocytes and play a non-redundant role in supporting clonal deletion and Treg selection^59,75,94,95^. In fact, recent studies have shown that presentation of AQP4 by thymic B cells is induced by CD40-mediated B cell licensing in the thymus and is required to establish T-cell tolerance to this self-antigen that is the target of autoimmune T-cell responses in neuromyelitis optica^96^. Here, we find that activated B cells are first detected between P3 and P7 and rise to become the predominant B cell subset at transitional ages. A recent study shows that B cell licensing also peaks in the perinatal period^75^. Consistent with our murine dataset, we find that the frequency of B cells increases in human thymuses across the perinatal transition. Together, these data suggest that thymic B cells at transitional ages play a conserved and critical role in establishing T cell tolerance to self-antigens expressed by germinal center B cells, with broader implications for tissue-specific self-tolerance.

Analysis of cDC subsets across the perinatal to juvenile transition also revealed dynamic changes in subset composition and transcriptional profiles of relevance to central tolerance. We find that aDC1s are most frequent within the thymic cDC compartment at perinatal (P0, P3) ages. Of the DC subsets, aDC1s most efficiently acquire and present mTEC-derived self-antigens, including Aire-dependent tissue restricted antigens (TRAs), on MHCI and MHCII^64,67^, and aDC1s have been shown to support Treg selection and clonal deletion^62,97^. Expression of *Aire* in the perinatal period is necessary and sufficient to support T-cell tolerance to peripheral tissues^84^, and *Aire*-dependent Treg selected in the perinatal thymus persist into adulthood and protect against autoimmunity in multiple tissues^13^. Taken together, these findings suggest that the high frequency of aDC1s in the perinatal period may play a critical role in selecting Treg that suppress autoimmunity to TRAs. Although the frequency of Tregs within the CD4SP compartment does not change across the transition, PD1 and CTLA4 expression are elevated on perinatal Treg, raising the possibility that this unique Treg subset may be selected by aDC1s and play an important role in suppressing autoimmune responses in tissues. cDC2s, which migrate into the thymus, increase in frequency during the transition (P7-P14), and we and others have shown that this subset is adept at Treg generation^72,74,98^. Because we find that transitional age P10 splenic Tregs are more efficient than adult Tregs at suppressing CD4+ T cell proliferation, the increased abundance of cDC2s during the transition suggests a potential role in selecting CD5^hi^GITR^hi^ Treg. However, because cDC2s continue to increase in abundance through the juvenile period, the unique transcriptional state of mTEC or DC subsets at the transition may control selection of highly suppressive Treg. To this point, it is notable that expression of genes associated with MHCII antigen processing and presentation were differentially expressed uniquely in transitional age DC subsets.

Recent studies have shown that IFN signaling induces activation of all thymic HAPC subsets^16,75^. Interestingly, we find a coordinated increase in genes associated with the type I IFN response starting at transitional ages not only in HAPC subsets, but also in TECs, mesenchymal and ECs. Increased IFN signaling across these diverse cell types is consistent with the finding that production of Type I and Type III IFNs in the thymus peaks at transitional ages^16^. Type I and Type III IFNs are expressed by mTECs in a partially *Aire*-dependent manner^17,99^. IFN signaling upregulates genes associated with antigen processing and presentation in the periphery^100^, which likely accounts for our finding that expression of MHCI antigen processing and presentation genes is temporally coordinated with enrichment of the Type I IFN signaling pathway in DCs during the transition. Consistent with this possibility, Type III IFN production by mTECs induces increased expression of MHCI by mTECs themselves^17^. The increase in antigen processing and presentation in DCs and TECs across the transition may contribute to the elevated levels of CD5 on post-positive selection thymocytes, as elevated MHC levels would induce stronger TCR signaling during thymocyte selection events. Furthermore, CD25+ Treg precursors, which are abundant in the perinatal thymus, require strong agonist signaling for selection^101^; thus, elevated antigen presentation on APCs during the transition could contribute to increased generation of these precursors.

Inverse age-associated changes in expression of type I IFN- and E2F-regulated genes occur in thymic epithelial, mesenchymal and endothelial cells as well as in hAPCs. That diverse cell types show similar temporal changes in expression of these genes indicates a widespread and coordinated shift in the transcriptional landscape throughout the thymic microenvironment. Additional studies are needed to determine if there is a cause-effect relationship between IFN- and E2F-induced transcriptional regulation. This may be the case as earlier studies have shown that type I IFN can inhibit E2F expression in tumor cell lines^102,103^. Regardless, the reciprocal transcriptional pattern of these gene sets suggests that the transition from perinatal thymus growth to juvenile homeostasis is accompanied by an increasingly pro-inflammatory state that likely influences thymocyte selection via enhanced antigen processing and presentation in TECs and hAPCs, which in turn augments the establishment of central tolerance.

In summary, this report provides high resolution data sets of previously unappreciated changes in the dynamic transcriptional profiles and composition of diverse thymic stromal cells and hAPCs across the perinatal to juvenile transition. We find these changes to be highly coordinated suggesting that global age-associated changes in the thymus microenvironment take place over the transition and may contribute to the distinct functional properties of T cells that develop in the perinatal versus adult thymus. These datasets offer the opportunity for further investigations into transcriptional programs regulating the switch between thymus growth and homeostasis. In addition, this resource will be useful for identifying additional age-associated changes in the cellular composition and gene expression profiles of TECs, mesenchymal cells, ECs and hAPCs that contribute to thymocyte development and establishment of central tolerance in the perinatal versus adult thymus.

## SUPPLEMENTAL INFORMATION

Document S1. Figures S1-S8

## ACKNOWLEDGEMENTS

The authors would like to thank Holly Stevenson and Jessica Podnar at the UT Austin Genome Sequencing and Analysis Facility for their assistance with generation of scRNAseq datasets. We also thank the veterinarians and staff of the animal facilities at our respective institutions. We thank Richard Salinas at the UT Austin’s Center for Biomedical Research Support and Pam Whitney in the department of Molecular Carcinogenesis and Epigenetics at the MD Anderson Cancer Center for technical assistance with flow cytometers. This research was supported by 1P01AI139449 from the NIAID/NIH to KL, LH, QL, NM, LE, and ER.

## AUTHOR CONTRIBUTIONS

Conceptualization: QL, NM, LE, ER Data Curation: BH, KL, QL

Formal analysis: AV, CM, AC, SK, BH, JS, SS, RZ, NS, SC, AM, LH

Funding Acquisition: KL, LH, QL, NM, LE, ER Methodology: QL, BH

Supervision: KL, LH, QL, NM, LE, ER

All authors wrote, edited, and reviewed the manuscript.

## STAR METHODS

### KEY RESOURCES TABLE

| REAGENT or RESOURCE | SOURCE | IDENTIFIER |
| --- | --- | --- |
| **Antibodies** |  |  |
| Anti-CD45-BV510; clone 30-F11 | Biolegend | Cat#103138; RRID:AB_2561392 |
| Anti-EPCAM-APC; clone G8.8 | Biolegend | Cat#118214; RRID:AB_1134102 |
| Anti-MHCII-APCcy7; clone M5/114.15.2 | Biolegend | Cat#107628; RRID:AB_2069377 |
| Anti-CD11c-PerCPcy5.5; clone N418 | Tonbo Biosciences | Cat#65-0114-U100; RRID:AB_2621886 |
| Ulex Europaeus Agglutinin I (UEA 1) | Vector Laboratories | Cat#FL-1061-2 |
| Anti-Ly51-Biotin; clone 6C3 | BD Biosciences | Cat#553159; RRID:AB_394671 |
| Anti-MHCII-PE-cy7; clone M5/114.15.2 | Biolegend | Cat#107629; RRID:AB_2069376 |
| Anti-CD45-AF700; clone 30-F11 | Biolegend | Cat#103127; RRID:AB_493714 |
| Anti-EpCAM-PE; clone G8.8 | Biolegend | Cat#118205; RRID:AB_1134172 |
| Anti-CD11c-Pacific Blue; clone N418 | Biolegend | Cat#117322; RRID:AB_755987 |
| Anti-Thy1.2-BV510; clone 53-2.1 | Biolegend | Cat#140319 |
| Anti-F4/80-APC/Cy7; clone BM8 | Biolegend | Cat#123117; RRID:AB_893477 |
| Anti-PDCA1-FITC; clone 927 | Biolegend | Cat#127007; RRID:AB_2028460 |
| Anti-Sirpa-APC; clone P84 | Biolegend | Cat#144013; RRID:AB_2564060 |
| Anti CD19-PerCP/Cy5.5; clone 1D3 | Biolegend | Cat#152405; RRID:AB_2629814 |
| Anti-CD11b-AF700; clone M1/70 | Biolegend | Cat#101222; RRID:AB_2738276 |
| Anti-CD11b-BUV395; clone M1/70 | BD Biosciences | Cat#563553; RRID:AB_2738276 |
| Anti-Sirpa-BV510; clone P84 | Biolegend | Cat#144032; RRID:AB_2810411 |
| Anti-PDCA1-PerCP/Cy5.5; clone 927 | Biolegend | Cat#127021; RRID:AB_2566646 |
| Anti-F4/80-BV785; clone BM8 | Biolegend | Cat#123141; RRID:AB_2563667 |
| Anti-CCR7-APC; clone 4B12 | Biolegend | Cat#120108; RRID:AB_389233 |
| Anti-CD63-AF700; clone NVG-2 | Biolegend | Cat#143924; RRID:AB_2832515 |
| Anti-XCR1-BV650; clone ZET | Biolegend | Cat#148220; RRID:AB_2566410 |
| Anti-CD3-Pacific Blue; clone 145-2C11 | Biolegend | Cat#100334; RRID:AB_2028475 |
| Anti-CD4-BV510; clone RM4-5 | Biolegend | Cat#100559; RRID:AB_2562608 |
| Anti-CD8-BV605; clone 53-6.7 | Biolegend | Cat#100744; RRID:<br>AB_2562609 |
| Anti-MHCI-PerCP/Cy5.5; clone AF6-88.5 | Biolegend | Cat#116516; RRID:<br>AB_1967133 |
| Anti-CD69-APC/Cy7; clone H1.2F3 | Biolegend | Cat#104526; RRID:<br>AB_10679041 |
| Anti-CD73-PE/Cy7; clone TY/11.8 | Biolegend | Cat#127224; RRID:<br>AB_2716103 |
| Anti-NK1.1 PE/Cy5; clone PK136 | Biolegend | Cat#108716; RRID:<br>AB_493590 |
| Anti-Gr1 PE/Cy5; clone RB6-8C5 | Biolegend | Cat#108410; RRID:<br>AB_313375 |
| Anti-CD19 PE/Cy5; clone 6D5 | Biolegend | Cat#115510; RRID:<br>AB_313645 |
| Anti-B220 PE/Cy5; clone RA3-6B2 | Biolegend | Cat#103210; RRID:<br>AB_312995 |
| Anti-CD11c PE/Cy5; clone N418 | Biolegend | Cat#117316; RRID:<br>AB_493566 |
| Anti-CD11b PE/Cy5; clone M1/70 | Biolegend | Cat#101210; RRID:<br>AB_312793 |
| Anti-TCRgd-PE/Cy5; clone eBioGL3 | Invitrogen | Cat#15-5711-82; RRID:<br>AB_468804 |
| Anti-Ter119-PE/Cy5; clone TER-119 | Biolegend | Cat#116210; RRID:<br>AB_313711 |
| Anti-CD25-AF488; clone PC61 | Biolegend | Cat#102017; RRID:<br>AB_493709 |
| Anti-CD73-PerCP/Cy5.5; clone TY/11.8 | Biolegend | Cat#127214; RRID:<br>AB_11219403 |
| Anti-CD44-AF700; clone IM7 | Biolegend | Cat#103026; RRID:<br>AB_493713 |
| Anti-PD1-APC; clone 29F.1A12 | Biolegend | Cat#135210; RRID:<br>AB_2159183 |
| Anti-GITR-PE/Cy7; clone YGITR765 | Biolegend | Cat#120222; RRID:<br>AB_528907 |
| Anti-Helios-APC; clone 22F6 | Biolegend | Cat#137221; RRID:<br>AB_10662535 |
| Anti-CD PE/Cy7; clone 53-7.3 | Biolegend | Cat#100622; RRID:<br>AB_2562773 |
| Anti-CTLA4-APC; clone UC10-4B9 | Biolegend | Cat#106310; RRID:<br>AB_2087653 |
| Anti-Icos-PE/Cy7; clone 7E.17G9 | Biolegend | Cat#117421; RRID:<br>AB_2860636 |
| Anti-FoxP3-PE; clone FJK-16s | ThermoFisher Scientific | Cat#12-5773-82; RRID:<br>AB_465936 |
| Anti-CD45-BV605; clone HI30 | Biolegend | Cat#304041; RRID:<br>AB_2562106 |
| Anti-EpCAM-APC; clone HEA-125 | Miltenyi Biotec | Cat#130-113-260 |
| Anti-CD11c-PE; clone 3.9 | Biolegend | Cat#301607; RRID:<br>AB_314175 |
| Anti-CD3-BV421; clone OKT3 | Biolegend | Cat#317343; RRID:<br>AB_2565848 |
| Anti-CD14-BV711; clone MΦP9 | BD Biosciences | Cat#563372; RRID:<br>AB_2744290 |
| Anti-MHCII-APC/Cy7; clone CVS20 | Novus Biologicals | Cat# NBP2-34848APCCY7 |
| UEA-1-FITC | ThermoFisher Scientific | Cat#L32476 |
| Anti-CD16-PE/Cy5; clone 3G8 | BD Biosciences | Cat#555408; RRID: AB_395808 |
| Anti-CD172a-PerCP-eFluor 710; clone 15-414 | ThermoFisher Scientific | Cat#46-1729-41/46-1729-42; RRID: AB_11043409 |
| Anti-CD317-PE/Cy7; clone RS38E | Biolegend | Cat#348416; RRID: AB_2716220 |
| Anti-CD7-BV786; clone M-T701 | BD Biosciences | Cat#740964; RRID: AB_2740589 |
| Anti-CD19-BV650; clone H1B19 | Biolegend | Cat#302238; RRID: AB_2562097 |
| Anti-Podoplanin-PE-Dazzle594; clone NC-08 | Biolegend | Cat#337027; RRID: AB_2750288 |
| Anti-human Vimentin | Leica Biosystems | Cat#NCL-VIM-V9 |
| Anti-cytokeratin 8; clone TROMA-1 | EMD Millipore | Cat#MABT329 |
| Anti-rabbit K5; clone Poly19055 | Biolegend | Cat#905501, RRID: AB_2565050 |
| Anti-chicken K14; clone Poly9060 | Biolegend | Cat#906004, RRID: AB_2616962 |
| Anti-cleaved caspase 3-FITC | Cell signaling | Cat#9669, RRID: AB_2069869 |
| Anti-Phospho-Rb (Ser807/811) XP<sup>R</sup> Rabbit mAb; Clone D20B12 | Cell Signaling | Cat# 8516 |
| Anti-β-Actin Rabbit mAb; clone D6A8 | Cell Signaling | Cat# 8457 |
| Anti-Rb antibody | GeneTex | Cat# GTX100545 |
| Anti-Rabbit HRP-Conjugated Goat Anti-Rabbit Ig(H+L) | Proteintech | Cat# SA00001-2 |
| **Biological samples** |  |  |
| Human thymus | Duke University Medical Center | N/A |
| **Chemicals, peptides, and recombinant proteins** |  |  |
| Propidium Iodide | Invitrogen<sup>TM</sup> | Cat#P3566 |
| Propidium Iodide | Enzo | Cat#ENZ-52403 |
| Streptavidin QDot605 | Invitrogen | Cat#Q10101MP |
| Fetal Bovine Serum | R&D systems | Cat#50-152-7066 |
| Fetal Bovine Serum | Sigma Aldrich | Cat#F2442 |
| EDTA | Boston Bioproducts Inc | Cat#BM-150 |
| EDTA | ThermoFisher Scientific | Cat# 15575020 |
| CD45 Microbeads, Mouse | Miltenyi Biotec | Cat#130-052-301 |
| LS columns | Miltenyi Biotec | Cat#130-042-401 |
| Zombie red Fixable Viability kit | Biolegend | Cat#423109; RRID: SCR_001134 |
| LIVE/DEAD<sup>TM</sup> Fixable Aqua Dead Cell stain kit | ThermoFisher Scientific | Cat#L34957 |
| Protease inhibitor cocktail | Millipore Sigma | Cat# 11836170001 |
| Phosphatase inhibitor cocktail | Millipore Sigma | Cat# P0044-1ML |
| Glycine | Fisher bioreagents | Cat#56-40-6 |
| Methanol | Fisher Scientific | Cat#CAS-67-56-1 |
| NaCl | Fisher Scientific | Cat#S271-3 |
| Tris base | Fisher Scientific | Cat#CAS-77-86-1 |
| Tween-20 | Millipore Sigma | Cat#X100 |
| 10XPBS, pH7.4 | Quality Biological | Cat# 119-069-101 |
| BSA | Serological Corporation | Cat# 3320-01 |
| Blocking reagent powder | Akoya Biosciences | Cat#FP1012 |
| PVDF membrane | Bio-Rad | Cat# 162-0181 |
| DAPI | Sigma Aldrich | Cat#D9542 |
| DNase I | Millipore Sigma | Cat#4716728001 |
| Collagenase/Dispase I | Millipore Sigma | Cat#10269638001 |
| Liberase<sup>TM</sup> Research grade | Millipore Sigma | Cat#5401127001 |
| 10X RBC lysis buffer | Biolegend | Cat#420301 |
| 1X ACK Lysing buffer | Lonza | Cat#BP10-548E |
| RPMI 1640 Medium (+25mM HEPES) | ThermoFisher Scientific | Cat#22400121 |
| Ham's F-12 Nutrient mix | Gibco<sup>TM</sup> | Cat#11765054 |
| Ultrapure<sup>TM</sup> BSA | Invitrogen<sup>TM</sup> | Cat#AM2616 |
| 10x PBS | Invitrogen<sup>TM</sup> | Cat#AM9624 |
| 0.5M EDTA | Invitrogen<sup>TM</sup> | Cat#AM9261 |
| CD3 microbeads, human | Miltenyi Biotec | Cat#130-097-043 |
| 4% Paraformaldehyde | ThermoFisher Scientific | Cat# J61899-AK |
| Trizol LS reagent | Invitrogen<sup>TM</sup> | Cat#15-596-018 |
| Linear acrylamide | Invitrogen<sup>TM</sup> | Cat#AM9520 |
| SuperSignal<sup>TM</sup> West Atto Ultimate Sensitivity Substrate | ThermoFisher Scientific | Cat#A38554 |
| Tissue-Tek<sup>®</sup> O.C.T. Compound | Sakura | Cat#4583 |
| Keratin 8 (TROMA-1) | EMD Millipore | Cat#MABT329 |
| ProLong<sup>TM</sup> Gold Antifade mountant | Invitrogen<sup>TM</sup> | Cat#P36930 |
| BOND epitope retrieval solution 1 | Leica Biosystems | Cat#AR9961 |
| DAB Enhancer | Leica Biosystems | Cat#AR9432 |
| Harris Hematoxylin nuclear stain | Lieca Biosystems | Cat#3801560 |
| Alexa Fluor<sup>®</sup> 488 AffiniPure<sup>TM</sup> Donkey Anti-Rat IgG (H+L) | Jackson Immunoresearch | Cat#712-546-153 |
| Alexa Fluor<sup>®</sup> 647 AffiniPure<sup>TM</sup> Donkey Anti-Rabbit IgG (H+L) | Jackson Immunoresearch | Cat#711-606-152 |
| Alexa Fluor<sup>TM</sup> 594 goat anti-chicken IgY (H+L) | Invitrogen<sup>TM</sup> | Cat#A11042 |
| **Critical commercial assays** |  |  |
| BD Cytofix/Cytoperm Fixation/Permeabilization kit | BD Biosciences | Cat#554714; RRID: SCR_013311 |
| Foxp3/Transcription Factor Staining Buffer kit | Tonbo Biosciences | Cat# TNB-0607-KIT |
| RNeasy Microprep kit | Qiagen | Cat#74104 |
| Superscript III First Strand cDNA Synthesis kit | Invitrogen<sup>TM</sup> | Cat#18080051 |
| Chromium Next GEM single-cell 3' reagent kit v3.1 | 10x Genomics | Cat#PN-1000268 |
| Bioanalyzer high sensitivity DNA kit | Agilent Technologies | Cat#5067-4626 |
| KAPA SYBR fast qPCR kit | Sigma Aldrich | Cat#SFUKB |
| **Deposited data** |  |  |
| Single-cell RNA-seq data-stromal cells | Kang et al., (revision submitted)<br>https://www.ncbi.nlm.nih.gov/geo/query/acc.cgi | GSE264402 |
| Single-cell RNA-seq data-stromal cells | Xiao and Kang et al.,<sup>104</sup> | GSE267785 |
| Single-cell RNA-seq data-TECs | This paper | GSE267926 |
| Single-cell RNA-seq data-hAPCs | This paper | GSE287039 |
| **Experimental Models: Organisms/strains** |  |  |
| C57BL/6J | The Jackson laboratory | Strain# 000664; RRID: IMSR_JAX:000664 |
| B6.Cg-Commd10Tg(Vav1-icre)A2Kio/J | The Jackson laboratory | Strain# 008610; RRID: IMSR_JAX:008610 |
| K5.cyclin D1 transgenic mice | Robles et al.,<sup>21</sup> | N/A |
| iLin28a transgenic mice | Yang et al.,<sup>20</sup> | N/A |
| **Oligonucleotides** |  |  |
| CCND1 | ThermoFisher scientific | Hs00765553_m1 |
| Lin28a | ThermoFisher scientific | Mm00524077_m1 |
| Mouse a-tubulin | ThermoFisher scientific | Mm00846967_g1 |
| Mouse GAPD Endogenous control | ThermoFisher scientific | 4352932E |
| **Software and algorithms** |  |  |
| Flowjo V10.10.0; V10.8.1; V10.9.0 | Treestar | https://www.flowjo.com/ |
| Prism V10.0.2, V10.0.3 | Graphpad | https://www.graphpad.com/ |
| Cell Ranger v6.0.1 | 10x Genomics | https://www.10xgenomics.com/support/software/cell-ranger/latest |
| Seurat v4.1.0 | Hao et al.,<sup>105</sup> Stuart et al.,<sup>106</sup> Butler et al.,<sup>107</sup> Satija et al.,<sup>108</sup> | https://satijalab.org/seurat/ |
| scMRMA v1.0 | Li et al.,<sup>109</sup> | N/A |
| ENCODE | The ENCODE Project Consortium<sup>110</sup>, Luo et al.,<sup>111</sup> | N/A |
| TEFA_Chip | Puente-Santamaria et al.,<sup>112</sup> | N/A |
| WebGestaltR | Wang and Liao et al<sup>113</sup> | https://cran.r-project.org/web/packages/WebGestaltR/index.html |
| rTENSOR | Li, Bien and Wells et al<sup>114</sup> | https://cran.r-project.org/web/packages/rTensor/index.html |
| R Version 4.1.0 | The R project | https://www.r-project.org/ |
| Fiji (Image J) version 2.14.0/1.54f | Schindelin et al.,<sup>115</sup> | RRID: SCR_002285 |
| **Other** |  |  |
| autoMACS Pro Separator | Miltenyi Biotec | N/A |
| LI-COR ODYSSEY Fc | LICORbio | N/A |
| Trans-blot System | Bio-Rad | Serial# 552BR027751 |
| BD LSRFortessa™ | BD (Becton, Dickinson and Company) | N/A |
| BD FACSAria™ Fusion | BD (Becton, Dickinson and Company) | N/A |
| BD FACSAria™ III | BD (Becton, Dickinson and Company) | N/A |
| Leica DMI8 microscope | Leica Biosystems | N/A |
| BOND-III stainer | Leica Biosystems | N/A |
| ABI 7900 HT fast real-time PCR system | Applied Biosystems | N/A |
| AB 7500 Real-time PCR system | Applied Biosystems | N/A |
| Nextseq 2000 sequencing system | Illumina | N/A |

### RESOURCE AVAILABILITY

#### Lead Contact

Further information and requests for resources and reagents should be directed to and will be fulfilled by the lead contact Ellen Richie

#### Materials availability

This study did not generate new unique reagents.

#### Data and code availability

Single-cell RNA-seq data from TECs, stromal cells and hAPCs are deposited at the Gene Expression Omnibus (GEO) and accession numbers are given in the key resources table. This paper does not report original code.

Any additional information required for reanalyzing data reported in this paper is available from the lead contact upon request.

### EXPERIMENTAL MODELS AND STUDY PARTICIPANT DETAILS

#### Mice

C57 BL6/J mice (RRID:IMSR_JAX:000664) and B6.Cg-Commd10Tg(Vav1-icre)A2Kio/J mice (RRID:IMSR_JAX:008610) were purchased from Jackson Laboratory and bred in-house. K5.Cyclin D1 mice were generated originally by C. Conti^21^ and maintained on a C57BL6/J background. *iLin28a* inducible transgenic mice were originally obtained from Dr. Eric Moss (Department of Molecular Biology, Rowan University, Stratford, NJ 08084 USA) and transferred from Dr. Jianfu Chen (Department of Genetics, University of Georgia) as previously reported^18,20^. All mice were maintained under specific pathogen-free conditions at The University of Texas at Austin, University of Texas MD Anderson Cancer Center, Smithville, and the University of Georgia. All experimental procedures were performed in accordance with the Institutional Animal Care and Use Committees.

### METHOD DETAILS

#### Cell isolation

Isolation of TECs for sorting and western blot was performed by cutting mouse thymi into small pieces and enzymatically digesting as previously described^116^. Briefly, cut thymic lobes were placed in conical tubes with 0.5ml (P0-P10) or 2ml of digestion buffer (P14-P28) containing 0.1mg/ml of Liberase TM (Sigma-Aldrich) and 20U/ml of DNAse I (Sigma-Aldrich) in PBS. Tubes were digested in a shaker incubator set to 37°C at 180rpm for 10 minutes for a total of 3 digestion rounds. At each round, supernatant was transferred into a conical tube containing FACS wash buffer (FWB) (1x PBS+2% Fetal Bovine Serum+5mM EDTA). Tubes were spun at 1500rpm for 5 minutes at 4°C and resuspended in 2ml of FACS buffer (P0-P7) or 5ml of FACS buffer (P10-P28). Single-cell suspension was filtered using a 70mm filter (Falcon) and then counted using a hemocytometer. For single-cell RNA sequencing, CD45+ thymocytes from digested thymi were depleted using the MACS Miltenyi mouse CD45 microbeads and LS columns (Miltenyi Biotec) according to manufacturer’s instructions. Briefly, 100ul of mouse CD45 beads were added to 1 x 10^8^ cells and incubated for 15mins on ice. following this, cell suspension was applied onto the LS columns and flow through containing CD45 depleted cells was collected. Cells were spun at 1500rpm for 5mins at 4°C processed for staining by flow cytometry. for Western blot, 1 x 10^8^ cells incubated with 100ul of mouse CD45 beads were depleted of CD45+ thymocytes using the autoMACS Pro Separator (Miltenyi Biotec) according to manufacturer’s instructions and processed for staining by flow cytometry.

For isolation of hAPCs, thymi were harvested and cut into small pieces for enzymatic digestion. They were placed in 50ml conical tubes containing 2ml of digestion buffer (0.1mg/ml of Liberase TM (Sigma-Aldrich) and 20U/ml of DNAse I (Sigma-Aldrich) in PBS). Thymi were digested for 12 minutes in a 37°C water bath each round for a total of 3 digestion rounds. At each digestion step, supernatant was collected and transferred to FACS tube containing 25ml of FACS wash buffer (PBS+2% Bovine calf serum+5mM EDTA). Cells were spun at 1250rpm for 5 mins at 4°C and processed for FACS analysis or sorting.

For isolation of thymocytes, thymuses were manually dissociated in FACS wash buffer and filtered through a 40mM cell strainer into PBS+2% bovine calf serum for flow cytometry staining.

Human thymus was obtained from anonymous patients undertaking corrective cardiac surgery where a portion of thymus was removed to expose the operative field. This tissue was not excised for the purpose of this study and would otherwise have been discarded. Age and sex of the donor were collected without any other identifying information. The use of anonymous discarded thymus tissue was approved by the Duke Institutional Review Board (Duke IRB Exempt Protocols #474 and #Pro00103028). Thymus tissues were stored overnight at 4°C in Ham’s F12 media (Invitrogen) 10% FBS (Sigma-Aldrich), washed in PBS, 1%BSA (Invitrogen) and then up to 5g of tissue was digested for 12 minutes at 37°C with occasional agitation in 20mL digestion buffer (PBS (Invitrogen), 0.52U/mL Liberase (Sigma-Aldrich) 20U/mL DNase I (Sigma-Aldrich). The released cells were added to 30mL wash buffer (PBS, 1% BSA, 5mM EDTA(Invitrogen) and the remaining tissue digested again in fresh digestion buffer. The digests were then pooled, pelleted, resuspended in wash buffer, filtered (70µm; Miltenyi Biotec), pelleted, and then resuspended in ACK lysis buffer (Lonza) for 1 minute for red blood cell lysis as needed. The cells were then rinsed in PBS and resuspended in wash buffer for counting.

Preparation of mouse thymic stromal cells for single-cell RNA sequencing is detailed in Kang et al, 2025 (revision submitted). Briefly, cut thymic lobes were digested in buffer containing 1640 RPMI (w/25mM HEPES) +2% FBS with 20μg/mL DNase I and 1mg/ml Collagenase/Dispase for a total of 3 rounds. Each round, thymi were incubated in a 37°C water bath for 20 mins to obtain a single-cell suspension. Following this, cells were centrifuged at 300rcf for 10mins at 4°C and incubated with 1X RBC lysis buffer (Biolegend) for 2 minutes. Cells were spun at 300rcf for 10mins at 4°C and resuspended in MACS buffer (filtered and degassed 0.5% BSA/2mM EDTA/1X PBS (w/o Ca2+, Mg2+)) for cell counting on the hemocytometer. CD45-cells for single-cell RNA sequencing were then collected by depletion of CD45-cells using anti-mouse CD45 microbeads according to manufacturer’s instructions (Miltenyi Biotec).

#### Flow cytometry and sorting

For quantification of TEC populations in WT and K5.D1 mice, 4 x 10^6^ cells were immunostained with the following anti-mouse fluorescent conjugated antibodies for 15mins at 4°C; CD45 (30-F11), EPCAM (G8.8), MHCII (M5/114.15.2) CD11c (N418), UEA, Ly51 (6C3), CD80 (16-10A1), SCA-1 (D7) and CD24 (M1/69).

Following staining, tubes were washed with FACS buffer at 1500rpm for 5mins at 4°C and resuspended in 600ul of FACS buffer for acquisition. For quantification of hAPC and DC subsets, 6 x 10^6^ cells were stained with the following anti-mouse antibodies; CD11b (M1/70), CD11c (N418), CD19 (1D3), EpCAM (G8.8), F4/80 (BM8), MHC II (M5/114.15.2), PDCA1 (927), Sirpa (P84), CCR7 (4B12), CD63 (NVG-2) and XCR1 (ZET). Cells were incubated for 40mins at room temperature and were washed at resuspended in FACS wash buffer +EDTA+propidium Iodide (Enzo). For quantification of thymocyte subsets, cells were stained with anti-mouse antibodies against the following cell surface markers; CD3 (145-2C11), CD4 (RM4-5), CD8 (53-6.7), MHCI (AF6-88.5), CD69 (H1.2F3), CD73 (TY/11.8), NK1.1 (PK136), Gr1 (RB6-8C5), CD19(6D5), B220(RA3-6B2), CD11c(N418), CD11b(M1/70), TCRγδ (eBioGL3), Ter119(TER-119), CD5(53-7.3). cells were incubated for 30mins at 4°C followed by centrifugation at 1400rpm for 5mins at 4°C. following surface staining, intracellular staining of cleaved caspase 3 was performed by fixing and permeabilizing cells using the BD cytofix/cytoperm Fixation/Permeabilization kit according to manufacturer’s instructions. Treg analysis was performed by surface staining of thymocytes using the following antibodies; CD3(145-2C11), CD4 (RM4-5), CD8 (53-6.7), CD25(PC61), CD73(TY/11.8), CD44(IM7), PD1(29F.1A12), GITR(YGITR765), Helios(22F6), CD5(53-7.3), CTLA4(UC10-4B9), Icos(7E.17G9) followed by intracellular staining of FOXP3(FJK-16s) using the FOXP3 TF staining buffer kit (tonbo Biosciences) according to manufacturer’s instructions. Viability for thymocyte subsets and Tregs was determined using Zombie Red™ Fixable Viability kit (Biolegend). Flow cytometry data was acquired on BD LSRFortessa^TM^ and analyzed using Flowjo software V10.9.0 or V10.10.0.

For quantification of human thymic antigen presenting cells, 10^8^ cells were labelled with 1μg/mL biotinylated anti-CD7 (124-1D1) in MACS buffer (PBS+0.5% FBS+2mM EDTA) for 15 minutes at 4°C, pelleted, and then incubated 15 minutes at 4°C for with magnetic beads coated with anti-CD3 and streptavidin (Miltenyi Biotec) in MACS buffer at the manufacturer’s recommended ratios. Cells were washed and resuspended in MACS buffer and then applied to a magnetic column (Miltenyi Biotec) to deplete CD3+ and CD7+ thymocytes. The eluted cells were then labelled for 40 minutes at room temperature with anti-human fluorescent labeled antibodies as follows; CD45 (HI30), CD326 (HEA-125), CD11c (3.9), CD3 (OKT3), CD14 (MΦP9), MHCII (CVS20), UEA-1, CD16 (3G8), CD172a (15-414), CD317 (RS38E), CD7 (M-T701), CD19 (HIB19) and podoplanin (NC-08) in PBS + 1%BSA, washed in PBS +1%BSA and stained with live/dead fix aqua (Invitrogen) in accordance with the manufacturer’s protocol, fixed in 4% PFA (ThermoFisher Scientific) and then analyzed on a BD LSRII flow cytometer. Antigen Presenting Cells were then gated and tabulated using FlowJo v10 according to the gating strategy shown in figure S5. Thymocyte depletion was confirmed by flow cytometry as shown.

Sorting of TECs for single-cell RNA sequencing was performed by immunostaining of CD45 depleted stromal cells with the following anti-mouse antibodies: CD45 (30-F11), EPCAM (G8.8), MHCII (M5/114.15.2) and CD11c (N418) and incubated for 15 minutes at 4°C. Cells were then washed and resuspended in FACS buffer for sorting on FACSAria^TM^ Fusion. EPCAM^+^CD45^-^ TECs were sorted and spun down in FACS buffer without EDTA. 3-8 x 10^5^ cells were submitted for single-cell RNA sequencing. For western blot, CD45 depleted stromal cells were stained with anti-mouse CD45 (30-F11) and EPCAM (G8.8) and incubated for 15 minutes at 4°C. Cells were then washed and resuspended in FACS buffer for sorting on FACSAria^TM^ III cell sorter.

Sorting of hAPCs for single-cell RNA sequencing was performed by immunostaining of enzymatically digested thymi with the following antibodies: MHCII (M5/114.15.2), CD45 (30-F11), EPCAM (G8.8), CD11c (N418), Thy1.2 (53-2.1), F4/80 (BM8), PDCA1 (927), Sirpa (P84) and CD19 (1D3) for 40mins at room temperature. Cells were resuspended in FACS wash buffer+ EDTA buffer plus propidium iodide (Enzo). Sorted cells were then spun down in FWB without EDTA, counted and resuspended at a concentration of 700-1200 cells/μL prior to submission to the sequencing core.

#### Western blot

TEC protein lysates were prepared in RIPA protein lysis buffer from pooled CD45^-^EpCam^+^ sorted cells at each age and each genotype. Each TEC sample was prepared from 5 to 8 mice. 2 *u*g TEC protein samples were subjected to western blotting by loading onto 5-15% gradient gels (Bio-RAD). After transfer, the PVDF membrane (Biorad) was probed with rabbit anti-pRB antibody (Cell Signaling Inc) for phospho-RB (pRB) as well as rabbit anti-Actin antibody (Cell Signaling Inc) as a loading control in the cold room for overnight incubation. After pRB blot, the PVDF membrane blot was subjected to mild stripping according to instructions (Abcam), then was blocked with 5% milk for an hour. Total RB (tRB) was detected with RB antibody (GeneTex) with an overnight incubation. Immunosignal was detected with SuperSignal West Atto Substrate (ThermoFisher Scientific). Western blotting band intensity was analyzed with LI-COR ODYSSEY Fc for pRB/Actin and tRB/Actin ratio.

#### qRT-PCR

qRT-PCR data of *Lin28a* is shown as previously reported^18^. Briefly, BM cells (T/B cell depleted) from 9-week-old Vav-iCre^+/-^;R26iLin28a and Vav-icre^-/-^;R26iLin28a mice were transferred into host *Foxn1^lacZ^* mutants (+/Z and Z/Z mice). Vav-iCre^+/-^;R26iLin28a and Vav-icre^-/-^;R26iLin28a BM progenitors recovered from host after 21 days was extracted using RNeasy micro kit (Qiagen) according to manufacturer’s instructions. qRT-PCR experiments were performed using the Lin28A Taqman Gene expression Assay kit (Mm00524077_m1) (ThermoFisher Scientific) and GAPDH FAM primer/probe as a control in a 10ml volume using the AB 7500 Sequence detector. qRT-PCR of *hCCND1* was performed using sorted EPCAM^+^CD45^-^TECs as described above from P3 and P28. EPCAM+CD45-TECs were sorted directly into Trizol (Ambion). mRNA from sorted TECs was extracted using chloroform and linear acrylamide (Ambion) according to manufacturer’s instructions. Following mRNA extraction, SuperScriptR III first strand cDNA synthesis kit (Invitrogen) was used to synthesize first strand-cDNA. qRT-PCR experiments were performed using the CCND1 Taqman gene expression assay kit (Hs00765553_m1) (ThermoFisher Scientific) and mouse α-tubulin (Mm00846967_g1) (ThermoFisher Scientific) was used as a control using the ABI 7900HT Fast Real-Time PCR system.

#### Immunofluorescence, Immunohistochemistry and microscopy

For immunofluorescence staining of Keratins, thymi harvested from mice of different ages were cleaned of connective tissues. Thymi were then placed in a mould containing OCT (Tissue-Tek) and flash frozen in methylbutane and dry ice. 10μm tissue sections were obtained on the cryostat. Thymuses were sectioned beginning to end to obtain sections at all depths. Sections representing the middle of the thymus were used for staining. Slides were left out at room temperature for 30mins to dry followed by fixation in acetone for 20mins at −20°C. slides were then washed in PBST (PBS+0.5% Tween 20) 3 times for 5 minutes each and blocked with TNB buffer for 15minutes at room temperature. Troma-1 supernatant was added to slides to stain overnight at 4°C. Next day, slides were washed in PBST (PBS+0.5% Tween 20) 3 times for 5 minutes each and Keratin 5 (Poly19055) and Keratin 14(poly9060) primary antibodies were added to stain for 2 hours at room temperature. Alternatively, Keratin 8 (TROMA-1), Keratin 5 (Poly19055) and Keratin 14(poly9060) primary antibodies were added to TNB blocked slides to stain for 2 hours at room temperature. Following washing in PBST (PBS+0.5% Tween 20) 3 times for 5 minutes each, secondary antibodies ((anti-rat 488, anti-rabbit 647 (Jackson Immunoresearch) and anti-chicken 594 (ThermoFisher)) diluted in TNB buffer were added to slides and incubated for 1hour at room temperature. slides were washed in PBST (PBS+0.5% Tween 20) 3 times for 5 minutes each and stained in DAPI (Sigma Aldrich) for 5 mins. Following washing in PBST 3 times 5 minutes each, slides were mounted with a coverslip using prolong gold (Invitrogen), imaged on Leica DMi8 microscope and analyzed using Fiji (ImageJ) software.

Human thymus was obtained from anonymous patients at Duke University Medical Center who were undergoing corrective cardiac surgery where a portion of the thymus was removed in order to expose the operative field. This tissue was not excised for the purpose of this study and would otherwise have been discarded. The age and sex of the donor were provided, without any other potentially identifying information. The use of anonymous discarded thymus tissue for this research was approved by the Duke University Institutional Review Board (Duke IRB Exempt Protocols #474 and #Pro00103028). Immunohistochemistry was performed on 4-5 µm formalin-fixed, paraffin-embedded sections of human thymus tissue using standard immunoperoxidase methodologies with a Leica Bond automated stainer, including antigen retrieval with ER1 (Leica; citrate with pH range of 5.9 to 6.1) for 20 minutes, vimentin antibody (clone V9; Leica), Bond polymer refine detection with 3, 3’-diaminobenzidine (brown) substrate (Leica), and a hematoxylin counterstain.

#### Library preparation and Single-cell RNA sequencing

Single-cell suspensions from TECs and hAPCs were processed in the University of Texas Genomic Sequencing and Analysis Facility. Cell suspensions were loaded on the Chromium Controller (10X Genomics) and processed for cDNA library generation following the manufacturer’s instructions for the Chromium Next GEM Single-cell 3’ Reagent Kit v3.1 (10X Genomics). The resulting libraries were examined for size and quality using the Bioanalyzer High Sensitivity DNA Kit (Agilent) and their concentrations were measured using the KAPA SYBR Fast qPCR kit (Roche). Samples were sequenced on the NovaSeq 6000 instrument (paired end, read 1: 28 cycles, read 2: 90 cycles) with a targeted depth of 50,000 reads/cell.

Library preparation and single-cell RNA sequencing of thymic stromal cells is provided in Kang et al, 2025 (revision submitted). Briefly, concentration of cells and viability were measured using a hemocytometer and countess 3. (ThermoFisher Scientific) following which,10,000 CD45-cells were loaded onto the Chromium controller and processed for cDNA library generation as given above. Samples were sequenced on Illumina NextSeq2000 P3 with 200 cycles.

#### Analysis of single-cell RNA sequencing data

Single-cell RNA-seq reads were processed using 10X Genomics Cell Ranger v6.0.1 to generate gene-by-cell count matrices^117^. Low-quality cells, defined as those with fewer than 500 expressed genes or with more than 10% of reads derived from mitochondrial genes, were excluded from downstream analyses. Following quality control, the single-cell RNA-seq data were normalized to 10,000 UMI counts, and the top 2,000 highly variable genes were identified by fitting the variance-mean relationship using Seurat v4.1.0^105–108^.The normalized data were scaled and principal component analysis (PCA) was performed for dimensionality reduction. The top 30 principal components were used to generate UMAP embeddings for visualization and to construct a k-nearest neighbor graph (k = 20). Louvain clustering was then applied to the graph using Seurat v4.1.0 to partition cells into non-overlapping clusters. Cell clusters were initially annotated using the marker-based scMRMA tool v1.0^109^, referencing markers reported in a previous study^31^. Annotations were further refined manually based on cluster-specific genes identified through differential expression analysis. Additionally, cells were assigned to G1, S, or G2M phases based on cell cycle scores calculated using Seurat v4.1.0.

#### Temporal expression pattern analysis

To investigate expression variation over time for each cell type, a pseudo-bulk gene expression profile was generated for each time point by averaging the expression levels of all cells from that cell type at the given time point. Principal component analysis (PCA) was performed on the pseudo-bulk profiles to better understand gene expression variability over time within the cell type of interest. Since the first principal component (PC1) typically captures temporal dynamics, and the second principal component (PC2) often reflects transitions, we sought to identify transcription factors (TFs) that might regulate these two processes. The top 200 genes contributing most to each principal component comprising the 100 genes with the highest positive contributions and 100 with the highest negative contributions were selected for further analysis. TF binding enrichment was performed on these gene sets using ChIP-seq data from the ENCODE database^110,111^ accessed via the TFEA.ChIP package^112^. Enrichment significance was assessed using a hypergeometric test, with TF-binding results meeting an FDR < 0.05 retained for further investigation.

To uncover gene clusters with shared temporal expression patterns, k-means clustering was applied to the pseudo-bulk profiles of the 1,000 most variable genes within the cell type of interest. Functional enrichment analysis was then conducted for each gene cluster using WebGestaltR, identifying enriched pathways at an FDR threshold of < 0.05. Hallmark and KEGG databases were used to identify Antigen processing and presentation pathway and type I IFN pathway genes enriched in k-means clustering.

#### Tensor decomposition

A three-dimensional tensor (gene×cell type×time) was constructed by combining the pseudo-bulk expression profiles at each time point for each cell type. The included cell types were cTEC, Ccl21^hi^ TEC, Aire^hi^ TEC, Aire^lo^ TEC, cDC1, cDC2, aDC1, aDC2, B cells, and macrophages. Endothelial and mesenchymal cells were excluded due to their limited number of time points. CP decomposition was performed on the tensor to extract a one-rank tensor for approximation using the rTensor package, and the percentage of the Frobenius norm (F-norm) explained by this approximation was reported. To assess the significance of the one-rank tensor, a background distribution of F-norm explained was generated by applying CP decomposition to tensors with the same number of genes, randomly sampled 1,000 times. The observed F-norm was then compared to the random sampling distribution to calculate statistical significance.

### QUANTIFICATION AND STATISTICAL ANALYSIS

Statistical comparisons between 2 groups were performed using multiple unpaired t-tests with FDR at 1% using two-stage step-up (Benjamini, Krieger, and Yekutieli) or using the Holm-Šídák method. Comparisons between 3 or more data groups was performed using one way ANOVA with Tukey’s or Dunnett’s multiple comparisons. *P<0.05, **P<0.01, ***P<0.001, ****P<0.0001. All calculations were performed using GraphPad Prism V10.0.2 or newer.

## REFERENCES

1. Haynes, B.F., Markert, M.L., Sempowski, G.D., Patel, D.D., and Hale, L.P. (2000). The role of the thymus in immune reconstitution in aging, bone marrow transplantation, and HIV-1 infection. Annu Rev Immunol 18, 529–560. 10.1146/annurev.immunol.18.1.529.

2. Steinmann, G.G., Klaus, B., and Muller-Hermelink, H.K. (1985). The involution of the ageing human thymic epithelium is independent of puberty. A morphometric study. Scand J Immunol 22, 563–575. 10.1111/j.1365-3083.1985.tb01916.x.

3. Chen, L., Xiao, S., and Manley, N.R. (2009). Foxn1 is required to maintain the postnatal thymic microenvironment in a dosage-sensitive manner. Blood 113, 567–574. 10.1182/blood-2008-05-156265.

4. Shah, D.K., and Zuniga-Pflucker, J.C. (2014). An overview of the intrathymic intricacies of T cell development. J Immunol 192, 4017–4023. 10.4049/jimmunol.1302259.

5. Ashby, K.M., and Hogquist, K.A. (2024). A guide to thymic selection of T cells. Nat Rev Immunol 24, 103–117. 10.1038/s41577-023-00911-8.

6. Hale, J.S., Boursalian, T.E., Turk, G.L., and Fink, P.J. (2006). Thymic output in aged mice. Proc Natl Acad Sci U S A 103, 8447–8452. 10.1073/pnas.0601040103.

7. Srinivasan, J., Lancaster, J.N., Singarapu, N., Hale, L.P., Ehrlich, L.I.R., and Richie, E.R. (2021). Age-Related Changes in Thymic Central Tolerance. Front Immunol 12, 676236. 10.3389/fimmu.2021.676236.

8. Tabilas, C., Smith, N.L., and Rudd, B.D. (2023). Shaping immunity for life: Layered development of CD8(+) T cells. Immunol Rev 315, 108–125. 10.1111/imr.13185.

9. Adkins, B., Leclerc, C., and Marshall-Clarke, S. (2004). Neonatal adaptive immunity comes of age. Nat Rev Immunol 4, 553–564. 10.1038/nri1394.

10. Havran, W.L., Carbone, A., and Allison, J.P. (1991). Murine T cells with invariant gamma delta antigen receptors: origin, repertoire, and specificity. Semin Immunol 3, 89–97.

11. Carding, S.R., and Egan, P.J. (2002). Gammadelta T cells: functional plasticity and heterogeneity. Nat Rev Immunol 2, 336–345. 10.1038/nri797.

12. Smith, N.L., Patel, R.K., Reynaldi, A., Grenier, J.K., Wang, J., Watson, N.B., Nzingha, K., Yee Mon, K.J., Peng, S.A., Grimson, A., et al. (2018). Developmental Origin Governs CD8(+) T Cell Fate Decisions during Infection. Cell 174, 117–130 e114. 10.1016/j.cell.2018.05.029.

13. Yang, S., Fujikado, N., Kolodin, D., Benoist, C., and Mathis, D. (2015). Immune tolerance. Regulatory T cells generated early in life play a distinct role in maintaining self-tolerance. Science 348, 589–594. 10.1126/science.aaa7017.

14. Ikuta, K., Kina, T., MacNeil, I., Uchida, N., Peault, B., Chien, Y.H., and Weissman, I.L. (1990). A developmental switch in thymic lymphocyte maturation potential occurs at the level of hematopoietic stem cells. Cell 62, 863–874. 10.1016/0092-8674(90)90262-d.

15. Notta, F., Zandi, S., Takayama, N., Dobson, S., Gan, O.I., Wilson, G., Kaufmann, K.B., McLeod, J., Laurenti, E., Dunant, C.F., et al. (2016). Distinct routes of lineage development reshape the human blood hierarchy across ontogeny. Science 351, aab2116. 10.1126/science.aab2116.

16. Ashby, K.M., Voboril, M., Salgado, O.C., Lee, S.T., Martinez, R.J., O’Connor, C.H., Breed, E.R., Xuan, S., Roll, C.R., Bachigari, S., et al. (2024). Sterile production of interferons in the thymus affects T cell repertoire selection. Sci Immunol 9, eadp1139. 10.1126/sciimmunol.adp1139.

17. Benhammadi, M., Mathe, J., Dumont-Lagace, M., Kobayashi, K.S., Gaboury, L., Brochu, S., and Perreault, C. (2020). IFN-lambda Enhances Constitutive Expression of MHC Class I Molecules on Thymic Epithelial Cells. J Immunol 205, 1268–1280. 10.4049/jimmunol.2000225.

18. Xiao, S., Zhang, W., and Manley, N.R. (2018). Thymic epithelial cell-derived signals control B progenitor formation and proliferation in the thymus by regulating Let-7 and Arid3a. PLoS One 13, e0193188. 10.1371/journal.pone.0193188.

19. Yuan, J., Nguyen, C.K., Liu, X., Kanellopoulou, C., and Muljo, S.A. (2012). Lin28b reprograms adult bone marrow hematopoietic progenitors to mediate fetal-like lymphopoiesis. Science 335, 1195–1200. 10.1126/science.1216557.

20. Yang, M., Yang, S.L., Herrlinger, S., Liang, C., Dzieciatkowska, M., Hansen, K.C., Desai, R., Nagy, A., Niswander, L., Moss, E.G., and Chen, J.F. (2015). Lin28 promotes the proliferative capacity of neural progenitor cells in brain development. Development 142, 1616–1627. 10.1242/dev.120543.

21. Robles, A.I., Larcher, F., Whalin, R.B., Murillas, R., Richie, E., Gimenez-Conti, I.B., Jorcano, J.L., and Conti, C.J. (1996). Expression of cyclin D1 in epithelial tissues of transgenic mice results in epidermal hyperproliferation and severe thymic hyperplasia. Proc Natl Acad Sci U S A 93, 7634–7638. 10.1073/pnas.93.15.7634.

22. Garfin, P.M., Min, D., Bryson, J.L., Serwold, T., Edris, B., Blackburn, C.C., Richie, E.R., Weinberg, K.I., Manley, N.R., Sage, J., and Viatour, P. (2013). Inactivation of the RB family prevents thymus involution and promotes thymic function by direct control of Foxn1 expression. J Exp Med 210, 1087–1097. 10.1084/jem.20121716.

23. Chinnam, M., and Goodrich, D.W. (2011). RB1, development, and cancer. Curr Top Dev Biol 94, 129–169. 10.1016/B978-0-12-380916-2.00005-X.

24. Iaquinta, P.J., and Lees, J.A. (2007). Life and death decisions by the E2F transcription factors. Curr Opin Cell Biol 19, 649–657. 10.1016/j.ceb.2007.10.006.

25. Burkhart, D.L., and Sage, J. (2008). Cellular mechanisms of tumour suppression by the retinoblastoma gene. Nat Rev Cancer 8, 671–682. 10.1038/nrc2399.

26. Dick, F.A., and Rubin, S.M. (2013). Molecular mechanisms underlying RB protein function. Nat Rev Mol Cell Biol 14, 297–306. 10.1038/nrm3567.

27. Topacio, B.R., Zatulovskiy, E., Cristea, S., Xie, S., Tambo, C.S., Rubin, S.M., Sage, J., Koivomagi, M., and Skotheim, J.M. (2019). Cyclin D-Cdk4,6 Drives Cell-Cycle Progression via the Retinoblastoma Protein’s C-Terminal Helix. Mol Cell 74, 758–770 e754. 10.1016/j.molcel.2019.03.020.

28. Sanidas, I., Morris, R., Fella, K.A., Rumde, P.H., Boukhali, M., Tai, E.C., Ting, D.T., Lawrence, M.S., Haas, W., and Dyson, N.J. (2019). A Code of Mono-phosphorylation Modulates the Function of RB. Mol Cell 73, 985–1000 e1006. 10.1016/j.molcel.2019.01.004.

29. Pryce, J.W., Bamber, A.R., Ashworth, M.T., Kiho, L., Malone, M., and Sebire, N.J. (2014). Reference ranges for organ weights of infants at autopsy: results of >1,000 consecutive cases from a single centre. BMC Clin Pathol 14, 18. 10.1186/1472-6890-14-18.

30. Nusser, A., Sagar, Swann, J.B., Krauth, B., Diekhoff, D., Calderon, L., Happe, C., Grun, D., and Boehm, T. (2022). Developmental dynamics of two bipotent thymic epithelial progenitor types. Nature 606, 165–171. 10.1038/s41586-022-04752-8.

31. Baran-Gale, J., Morgan, M.D., Maio, S., Dhalla, F., Calvo-Asensio, I., Deadman, M.E., Handel, A.E., Maynard, A., Chen, S., Green, F., et al. (2020). Ageing compromises mouse thymus function and remodels epithelial cell differentiation. Elife 9. 10.7554/eLife.56221.

32. Wells, K.L., Miller, C.N., Gschwind, A.R., Wei, W., Phipps, J.D., Anderson, M.S., and Steinmetz, L.M. (2020). Combined transient ablation and single-cell RNA-sequencing reveals the development of medullary thymic epithelial cells. Elife 9. 10.7554/eLife.60188.

33. Bornstein, C., Nevo, S., Giladi, A., Kadouri, N., Pouzolles, M., Gerbe, F., David, E., Machado, A., Chuprin, A., Toth, B., et al. (2018). Single-cell mapping of the thymic stroma identifies IL-25-producing tuft epithelial cells. Nature 559, 622–626. 10.1038/s41586-018-0346-1.

34. Kernfeld, E.M., Genga, R.M.J., Neherin, K., Magaletta, M.E., Xu, P., and Maehr, R. (2018). A Single-Cell Transcriptomic Atlas of Thymus Organogenesis Resolves Cell Types and Developmental Maturation. Immunity 48, 1258–1270 e1256. 10.1016/j.immuni.2018.04.015.

35. Kousa, A.I., Jahn, L., Zhao, K., Flores, A.E., Acenas, D., 2nd, Lederer, E., Argyropoulos, K.V., Lemarquis, A.L., Granadier, D., Cooper, K., et al. (2024). Age-related epithelial defects limit thymic function and regeneration. Nat Immunol. 10.1038/s41590-024-01915-9.

36. Miragaia, R.J., Zhang, X., Gomes, T., Svensson, V., Ilicic, T., Henriksson, J., Kar, G., and Lonnberg, T. (2018). Single-cell RNA-sequencing resolves self-antigen expression during mTEC development. Sci Rep 8, 685. 10.1038/s41598-017-19100-4.

37. Ohigashi, I., White, A.J., Yang, M.T., Fujimori, S., Tanaka, Y., Jacques, A., Kiyonari, H., Matsushita, Y., Turan, S., Kelly, M.C., et al. (2024). Developmental conversion of thymocyte-attracting cells into self-antigen-displaying cells in embryonic thymus medulla epithelium. Elife 12. 10.7554/eLife.92552.

38. Miyao, T., Miyauchi, M., Kelly, S.T., Terooatea, T.W., Ishikawa, T., Oh, E., Hirai, S., Horie, K., Takakura, Y., Ohki, H., et al. (2022). Integrative analysis of scRNA-seq and scATAC-seq revealed transit-amplifying thymic epithelial cells expressing autoimmune regulator. Elife 11. 10.7554/eLife.73998.

39. Venables, T., Griffith, A.V., DeAraujo, A., and Petrie, H.T. (2019). Dynamic changes in epithelial cell morphology control thymic organ size during atrophy and regeneration. Nat Commun 10, 4402. 10.1038/s41467-019-11879-2.

40. Nakagawa, Y., Ohigashi, I., Nitta, T., Sakata, M., Tanaka, K., Murata, S., Kanagawa, O., and Takahama, Y. (2012). Thymic nurse cells provide microenvironment for secondary T cell receptor alpha rearrangement in cortical thymocytes. Proc Natl Acad Sci U S A 109, 20572–20577. 10.1073/pnas.1213069109.

41. Michelson, D.A., Hase, K., Kaisho, T., Benoist, C., and Mathis, D. (2022). Thymic epithelial cells co-opt lineage-defining transcription factors to eliminate autoreactive T cells. Cell 185, 2542–2558 e2518. 10.1016/j.cell.2022.05.018.

42. Givony, T., Leshkowitz, D., Del Castillo, D., Nevo, S., Kadouri, N., Dassa, B., Gruper, Y., Khalaila, R., Ben-Nun, O., Gome, T., et al. (2023). Author Correction: Thymic mimetic cells function beyond self-tolerance. Nature 624, E4. 10.1038/s41586-023-06881-0.

43. Subramanian, A., Tamayo, P., Mootha, V.K., Mukherjee, S., Ebert, B.L., Gillette, M.A., Paulovich, A., Pomeroy, S.L., Golub, T.R., Lander, E.S., and Mesirov, J.P. (2005). Gene set enrichment analysis: a knowledge-based approach for interpreting genome-wide expression profiles. Proc Natl Acad Sci U S A 102, 15545–15550. 10.1073/pnas.0506580102.

44. Mootha, V.K., Lindgren, C.M., Eriksson, K.F., Subramanian, A., Sihag, S., Lehar, J., Puigserver, P., Carlsson, E., Ridderstrale, M., Laurila, E., et al. (2003). PGC-1alpha-responsive genes involved in oxidative phosphorylation are coordinately downregulated in human diabetes. Nat Genet 34, 267–273. 10.1038/ng1180.

45. Jeremiah, N., Ferran, H., Antoniadou, K., De Azevedo, K., Nikolic, J., Maurin, M., Benaroch, P., and Manel, N. (2023). RELA tunes innate-like interferon I/III responses in human T cells. J Exp Med 220. 10.1084/jem.20220666.

46. Kochupurakkal, B.S., Wang, Z.C., Hua, T., Culhane, A.C., Rodig, S.J., Rajkovic-Molek, K., Lazaro, J.B., Richardson, A.L., Biswas, D.K., and Iglehart, J.D. (2015). RelA-Induced Interferon Response Negatively Regulates Proliferation. PLoS One 10, e0140243. 10.1371/journal.pone.0140243.

47. Xing, Y., Wang, X., Jameson, S.C., and Hogquist, K.A. (2016). Late stages of T cell maturation in the thymus involve NF-kappaB and tonic type I interferon signaling. Nat Immunol 17, 565–573. 10.1038/ni.3419.

48. Larouche, J.D., Laumont, C.M., Trofimov, A., Vincent, K., Hesnard, L., Brochu, S., Cote, C., Humeau, J.F., Bonneil, E., Lanoix, J., et al. (2024). Transposable elements regulate thymus development and function. Elife 12. 10.7554/eLife.91037.

49. Klug, D.B., Carter, C., Crouch, E., Roop, D., Conti, C.J., and Richie, E.R. (1998). Interdependence of cortical thymic epithelial cell differentiation and T-lineage commitment. Proc Natl Acad Sci U S A 95, 11822–11827. 10.1073/pnas.95.20.11822.

50. Manley, N.R., Richie, E.R., Blackburn, C.C., Condie, B.G., and Sage, J. (2011). Structure and function of the thymic microenvironment. Front Biosci (Landmark Ed) 16, 2461–2477. 10.2741/3866.

51. Nitta, T., and Takayanagi, H. (2020). Non-Epithelial Thymic Stromal Cells: Unsung Heroes in Thymus Organogenesis and T Cell Development. Front Immunol 11, 620894. 10.3389/fimmu.2020.620894.

52. James, K.D., Jenkinson, W.E., and Anderson, G. (2021). Non-Epithelial Stromal Cells in Thymus Development and Function. Front Immunol 12, 634367. 10.3389/fimmu.2021.634367.

53. Cuddihy, A.R., Ge, S., Zhu, J., Jang, J., Chidgey, A., Thurston, G., Boyd, R., and Crooks, G.M. (2009). VEGF-mediated cross-talk within the neonatal murine thymus. Blood 113, 2723–2731. 10.1182/blood-2008-06-162040.

54. Huang, H., Wang, Z., Zhang, Y., Pradhan, R.N., Ganguly, D., Chandra, R., Murimwa, G., Wright, S., Gu, X., Maddipati, R., et al. (2022). Mesothelial cell-derived antigen-presenting cancer-associated fibroblasts induce expansion of regulatory T cells in pancreatic cancer. Cancer Cell 40, 656–673 e657. 10.1016/j.ccell.2022.04.011.

55. Nitta, T., Ota, A., Iguchi, T., Muro, R., and Takayanagi, H. (2021). The fibroblast: An emerging key player in thymic T cell selection. Immunol Rev 302, 68–85. 10.1111/imr.12985.

56. Nitta, T., Tsutsumi, M., Nitta, S., Muro, R., Suzuki, E.C., Nakano, K., Tomofuji, Y., Sawa, S., Okamura, T., Penninger, J.M., and Takayanagi, H. (2020). Fibroblasts as a source of self-antigens for central immune tolerance. Nat Immunol 21, 1172–1180. 10.1038/s41590-020-0756-8.

57. Lei, Y., Ripen, A.M., Ishimaru, N., Ohigashi, I., Nagasawa, T., Jeker, L.T., Bosl, M.R., Hollander, G.A., Hayashi, Y., Malefyt Rde, W., et al. (2011). Aire-dependent production of XCL1 mediates medullary accumulation of thymic dendritic cells and contributes to regulatory T cell development. J Exp Med 208, 383–394. 10.1084/jem.20102327.

58. Klein, L., and Petrozziello, E. (2024). Antigen presentation for central tolerance induction. Nat Rev Immunol. 10.1038/s41577-024-01076-8.

59. Yamano, T., Nedjic, J., Hinterberger, M., Steinert, M., Koser, S., Pinto, S., Gerdes, N., Lutgens, E., Ishimaru, N., Busslinger, M., et al. (2015). Thymic B Cells Are Licensed to Present Self Antigens for Central T Cell Tolerance Induction. Immunity 42, 1048–1061. 10.1016/j.immuni.2015.05.013.

60. Bonasio, R., Scimone, M.L., Schaerli, P., Grabie, N., Lichtman, A.H., and von Andrian, U.H. (2006). Clonal deletion of thymocytes by circulating dendritic cells homing to the thymus. Nat Immunol 7, 1092–1100. 10.1038/ni1385.

61. Atibalentja, D.F., Murphy, K.M., and Unanue, E.R. (2011). Functional redundancy between thymic CD8alpha+ and Sirpalpha+ conventional dendritic cells in presentation of blood-derived lysozyme by MHC class II proteins. J Immunol 186, 1421–1431. 10.4049/jimmunol.1002587.

62. Perry, J.S.A., Russler-Germain, E.V., Zhou, Y.W., Purtha, W., Cooper, M.L., Choi, J., Schroeder, M.A., Salazar, V., Egawa, T., Lee, B.C., et al. (2018). Transfer of Cell-Surface Antigens by Scavenger Receptor CD36 Promotes Thymic Regulatory T Cell Receptor Repertoire Development and Allo-tolerance. Immunity 48, 1271. 10.1016/j.immuni.2018.05.011.

63. Kroger, C.J., Spidale, N.A., Wang, B., and Tisch, R. (2017). Thymic Dendritic Cell Subsets Display Distinct Efficiencies and Mechanisms of Intercellular MHC Transfer. J Immunol 198, 249–256. 10.4049/jimmunol.1601516.

64. Voboril, M., Brezina, J., Brabec, T., Dobes, J., Ballek, O., Dobesova, M., Manning, J., Blumberg, R.S., and Filipp, D. (2022). A model of preferential pairing between epithelial and dendritic cells in thymic antigen transfer. Elife 11. 10.7554/eLife.71578.

65. Hadeiba, H., Lahl, K., Edalati, A., Oderup, C., Habtezion, A., Pachynski, R., Nguyen, L., Ghodsi, A., Adler, S., and Butcher, E.C. (2012). Plasmacytoid dendritic cells transport peripheral antigens to the thymus to promote central tolerance. Immunity 36, 438–450. 10.1016/j.immuni.2012.01.017.

66. Breed, E.R., Voboril, M., Ashby, K.M., Martinez, R.J., Qian, L., Wang, H., Salgado, O.C., O’Connor, C.H., and Hogquist, K.A. (2022). Type 2 cytokines in the thymus activate Sirpalpha(+) dendritic cells to promote clonal deletion. Nat Immunol 23, 1042–1051. 10.1038/s41590-022-01218-x.

67. Srinivasan J, H.B., Yang Y, Moore C, Moore JF, Calindi A, Selden HJ, Heiser CN, Liu Q, Lau KS, Ehrlich LIR. (2023). Single-cell transcriptomics reveals heterogenous thymic dendritic cell subsets with distinct functions and requirements for thymocyte-regulated crosstalk. bioRxiv. 10.1101/2023.12.18.572281.

68. Oh, J., Wu, N., Barczak, A.J., Barbeau, R., Erle, D.J., and Shin, J.S. (2018). CD40 Mediates Maturation of Thymic Dendritic Cells Driven by Self-Reactive CD4(+) Thymocytes and Supports Development of Natural Regulatory T Cells. J Immunol 200, 1399–1412. 10.4049/jimmunol.1700768.

69. Ardouin, L., Luche, H., Chelbi, R., Carpentier, S., Shawket, A., Montanana Sanchis, F., Santa Maria, C., Grenot, P., Alexandre, Y., Gregoire, C., et al. (2016). Broad and Largely Concordant Molecular Changes Characterize Tolerogenic and Immunogenic Dendritic Cell Maturation in Thymus and Periphery. Immunity 45, 305–318. 10.1016/j.immuni.2016.07.019.

70. Goldschneider, I., and Cone, R.E. (2003). A central role for peripheral dendritic cells in the induction of acquired thymic tolerance. Trends Immunol 24, 77–81. 10.1016/s1471-4906(02)00038-8.

71. Vollmann, E.H., Rattay, K., Barreiro, O., Thiriot, A., Fuhlbrigge, R.A., Vrbanac, V., Kim, K.W., Jung, S., Tager, A.M., and von Andrian, U.H. (2021). Specialized transendothelial dendritic cells mediate thymic T-cell selection against blood-borne macromolecules. Nat Commun 12, 6230. 10.1038/s41467-021-26446-x.

72. Proietto, A.I., van Dommelen, S., Zhou, P., Rizzitelli, A., D’Amico, A., Steptoe, R.J., Naik, S.H., Lahoud, M.H., Liu, Y., Zheng, P., et al. (2008). Dendritic cells in the thymus contribute to T-regulatory cell induction. Proc Natl Acad Sci U S A 105, 19869–19874. 10.1073/pnas.0810268105.

73. Zegarra-Ruiz, D.F., Kim, D.V., Norwood, K., Kim, M., Wu, W.H., Saldana-Morales, F.B., Hill, A.A., Majumdar, S., Orozco, S., Bell, R., et al. (2021). Thymic development of gut-microbiota-specific T cells. Nature 594, 413–417. 10.1038/s41586-021-03531-1.

74. Hu, Z., Li, Y., Van Nieuwenhuijze, A., Selden, H.J., Jarrett, A.M., Sorace, A.G., Yankeelov, T.E., Liston, A., and Ehrlich, L.I.R. (2017). CCR7 Modulates the Generation of Thymic Regulatory T Cells by Altering the Composition of the Thymic Dendritic Cell Compartment. Cell Rep 21, 168–180. 10.1016/j.celrep.2017.09.016.

75. Martinez, R.J., Breed, E.R., Worota, Y., Ashby, K.M., Voboril, M., Mathes, T., Salgado, O.C., O’Connor, C.H., Kotenko, S.V., and Hogquist, K.A. (2023). Type III interferon drives thymic B cell activation and regulatory T cell generation. Proc Natl Acad Sci U S A 120, e2220120120. 10.1073/pnas.2220120120.

76. Lazear, H.M., Schoggins, J.W., and Diamond, M.S. (2019). Shared and Distinct Functions of Type I and Type III Interferons. Immunity 50, 907–923. 10.1016/j.immuni.2019.03.025.

77. Perera, J., Meng, L., Meng, F., and Huang, H. (2013). Autoreactive thymic B cells are efficient antigen-presenting cells of cognate self-antigens for T cell negative selection. Proc Natl Acad Sci U S A 110, 17011–17016. 10.1073/pnas.1313001110.

78. Akashi, K., Richie, L.I., Miyamoto, T., Carr, W.H., and Weissman, I.L. (2000). B lymphopoiesis in the thymus. J Immunol 164, 5221–5226. 10.4049/jimmunol.164.10.5221.

79. Dong, M., Artusa, P., Kelly, S.A., Fournier, M., Baldwin, T.A., Mandl, J.N., and Melichar, H.J. (2017). Alterations in the Thymic Selection Threshold Skew the Self-Reactivity of the TCR Repertoire in Neonates. J Immunol 199, 965–973. 10.4049/jimmunol.1602137.

80. Azzam, H.S., Grinberg, A., Lui, K., Shen, H., Shores, E.W., and Love, P.E. (1998). CD5 expression is developmentally regulated by T cell receptor (TCR) signals and TCR avidity. J Exp Med 188, 2301–2311. 10.1084/jem.188.12.2301.

81. Mandl, J.N., Monteiro, J.P., Vrisekoop, N., and Germain, R.N. (2013). T cell-positive selection uses self-ligand binding strength to optimize repertoire recognition of foreign antigens. Immunity 38, 263–274. 10.1016/j.immuni.2012.09.011.

82. McCaughtry, T.M., Wilken, M.S., and Hogquist, K.A. (2007). Thymic emigration revisited. J Exp Med 204, 2513–2520. 10.1084/jem.20070601.

83. Sawicka, M., Stritesky, G.L., Reynolds, J., Abourashchi, N., Lythe, G., Molina-Paris, C., and Hogquist, K.A. (2014). From pre-DP, post-DP, SP4, and SP8 Thymocyte Cell Counts to a Dynamical Model of Cortical and Medullary Selection. Front Immunol 5, 19. 10.3389/fimmu.2014.00019.

84. Guerau-de-Arellano, M., Martinic, M., Benoist, C., and Mathis, D. (2009). Neonatal tolerance revisited: a perinatal window for Aire control of autoimmunity. J Exp Med 206, 1245–1252. 10.1084/jem.20090300.

85. Tai, X., Van Laethem, F., Pobezinsky, L., Guinter, T., Sharrow, S.O., Adams, A., Granger, L., Kruhlak, M., Lindsten, T., Thompson, C.B., et al. (2012). Basis of CTLA-4 function in regulatory and conventional CD4(+) T cells. Blood 119, 5155–5163. 10.1182/blood-2011-11-388918.

86. Wyss, L., Stadinski, B.D., King, C.G., Schallenberg, S., McCarthy, N.I., Lee, J.Y., Kretschmer, K., Terracciano, L.M., Anderson, G., Surh, C.D., et al. (2016). Affinity for self antigen selects Treg cells with distinct functional properties. Nat Immunol 17, 1093–1101. 10.1038/ni.3522.

87. Mahmud, S.A., Manlove, L.S., Schmitz, H.M., Xing, Y., Wang, Y., Owen, D.L., Schenkel, J.M., Boomer, J.S., Green, J.M., Yagita, H., et al. (2014). Costimulation via the tumor-necrosis factor receptor superfamily couples TCR signal strength to the thymic differentiation of regulatory T cells. Nat Immunol 15, 473–481. 10.1038/ni.2849.

88. Anderson, G., and Takahama, Y. (2012). Thymic epithelial cells: working class heroes for T cell development and repertoire selection. Trends Immunol 33, 256–263. 10.1016/j.it.2012.03.005.

89. Boehm, T. (2008). Thymus development and function. Curr Opin Immunol 20, 178–184. 10.1016/j.coi.2008.03.001.

90. Bryson, J.L., Griffith, A.V., Hughes, B., 3rd, Saito, F., Takahama, Y., Richie, E.R., and Manley, N.R. (2013). Cell-autonomous defects in thymic epithelial cells disrupt endothelial-perivascular cell interactions in the mouse thymus. PLoS One 8, e65196. 10.1371/journal.pone.0065196.

91. Wertheimer, T., Velardi, E., Tsai, J., Cooper, K., Xiao, S., Kloss, C.C., Ottmuller, K.J., Mokhtari, Z., Brede, C., deRoos, P., et al. (2018). Production of BMP4 by endothelial cells is crucial for endogenous thymic regeneration. Sci Immunol 3. 10.1126/sciimmunol.aal2736.

92. Hartmann, W., Koch, A., Brune, H., Waha, A., Schuller, U., Dani, I., Denkhaus, D., Langmann, W., Bode, U., Wiestler, O.D., et al. (2005). Insulin-like growth factor II is involved in the proliferation control of medulloblastoma and its cerebellar precursor cells. Am J Pathol 166, 1153–1162. 10.1016/S0002-9440(10)62335-8.

93. Preusser, M., De Mattos-Arruda, L., Thill, M., Criscitiello, C., Bartsch, R., Ruhstaller, T., de Azambuja, E., and Zielinski, C.C. (2018). CDK4/6 inhibitors in the treatment of patients with breast cancer: summary of a multidisciplinary round-table discussion. ESMO Open 3, e000368. 10.1136/esmoopen-2018-000368.

94. Lu, F.T., Yang, W., Wang, Y.H., Ma, H.D., Tang, W., Yang, J.B., Li, L., Ansari, A.A., and Lian, Z.X. (2015). Thymic B cells promote thymus-derived regulatory T cell development and proliferation. J Autoimmun 61, 62–72. 10.1016/j.jaut.2015.05.008.

95. Walters, S.N., Webster, K.E., Daley, S., and Grey, S.T. (2014). A role for intrathymic B cells in the generation of natural regulatory T cells. J Immunol 193, 170–176. 10.4049/jimmunol.1302519.

96. Afzali, A.M., Nirschl, L., Sie, C., Pfaller, M., Ulianov, O., Hassler, T., Federle, C., Petrozziello, E., Kalluri, S.R., Chen, H.H., et al. (2024). B cells orchestrate tolerance to the neuromyelitis optica autoantigen AQP4. Nature 627, 407–415. 10.1038/s41586-024-07079-8.

97. Perry, J.S.A., Lio, C.J., Kau, A.L., Nutsch, K., Yang, Z., Gordon, J.I., Murphy, K.M., and Hsieh, C.S. (2014). Distinct contributions of Aire and antigen-presenting-cell subsets to the generation of self-tolerance in the thymus. Immunity 41, 414–426. 10.1016/j.immuni.2014.08.007.

98. Herbin, O., Bonito, A.J., Jeong, S., Weinstein, E.G., Rahman, A.H., Xiong, H., Merad, M., and Alexandropoulos, K. (2016). Medullary thymic epithelial cells and CD8alpha(+) dendritic cells coordinately regulate central tolerance but CD8alpha(+) cells are dispensable for thymic regulatory T cell production. J Autoimmun 75, 141–149. 10.1016/j.jaut.2016.08.002.

99. Lienenklaus, S., Cornitescu, M., Zietara, N., Lyszkiewicz, M., Gekara, N., Jablonska, J., Edenhofer, F., Rajewsky, K., Bruder, D., Hafner, M., et al. (2009). Novel reporter mouse reveals constitutive and inflammatory expression of IFN-beta in vivo. J Immunol 183, 3229–3236. 10.4049/jimmunol.0804277.

100. McNab, F., Mayer-Barber, K., Sher, A., Wack, A., and O’Garra, A. (2015). Type I interferons in infectious disease. Nat Rev Immunol 15, 87–103. 10.1038/nri3787.

101. Owen, D.L., Mahmud, S.A., Sjaastad, L.E., Williams, J.B., Spanier, J.A., Simeonov, D.R., Ruscher, R., Huang, W., Proekt, I., Miller, C.N., et al. (2019). Thymic regulatory T cells arise via two distinct developmental programs. Nat Immunol 20, 195–205. 10.1038/s41590-018-0289-6.

102. Iwase, S., Furukawa, Y., Kikuchi, J., Nagai, M., Terui, Y., Nakamura, M., and Yamada, H. (1997). Modulation of E2F activity is linked to interferon-induced growth suppression of hematopoietic cells. J Biol Chem 272, 12406–12414. 10.1074/jbc.272.19.12406.

103. Furukawa, Y., Iwase, S., Kikuchi, J., Nakamura, M., Yamada, H., and Matsuda, M. (1999). Transcriptional repression of the E2F-1 gene by interferon-alpha is mediated through induction of E2F-4/pRB and E2F-4/p130 complexes. Oncogene 18, 2003–2014. 10.1038/sj.onc.1202500.

104. Xiao, S., Kang, S.W., Oliva, K.E., Zhang, W., Klonowski, K.D., and Manley, N.R. (2024). Premature thymic involution in young Foxn1lacz mutant mice causes peripheral T cell phenotypes similar to aging-induced immunosenescence. bioRxiv. 10.1101/2024.04.23.590170.

105. Hao, Y., Hao, S., Andersen-Nissen, E., Mauck, W.M., 3rd, Zheng, S., Butler, A., Lee, M.J., Wilk, A.J., Darby, C., Zager, M., et al. (2021). Integrated analysis of multimodal single-cell data. Cell 184, 3573–3587 e3529. 10.1016/j.cell.2021.04.048.

106. Stuart, T., Butler, A., Hoffman, P., Hafemeister, C., Papalexi, E., Mauck, W.M., 3rd, Hao, Y., Stoeckius, M., Smibert, P., and Satija, R. (2019). Comprehensive Integration of Single-Cell Data. Cell 177, 1888–1902 e1821. 10.1016/j.cell.2019.05.031.

107. Butler, A., Hoffman, P., Smibert, P., Papalexi, E., and Satija, R. (2018). Integrating single-cell transcriptomic data across different conditions, technologies, and species. Nat Biotechnol 36, 411–420. 10.1038/nbt.4096.

108. Satija, R., Farrell, J.A., Gennert, D., Schier, A.F., and Regev, A. (2015). Spatial reconstruction of single-cell gene expression data. Nat Biotechnol 33, 495–502. 10.1038/nbt.3192.

109. Li, J., Sheng, Q., Shyr, Y., and Liu, Q. (2022). scMRMA: single cell multiresolution marker-based annotation. Nucleic Acids Res 50, e7. 10.1093/nar/gkab931.

110. Consortium, E.P. (2012). An integrated encyclopedia of DNA elements in the human genome. Nature 489, 57–74. 10.1038/nature11247.

111. Luo, Y., Hitz, B.C., Gabdank, I., Hilton, J.A., Kagda, M.S., Lam, B., Myers, Z., Sud, P., Jou, J., Lin, K., et al. (2020). New developments on the Encyclopedia of DNA Elements (ENCODE) data portal. Nucleic Acids Res 48, D882–D889. 10.1093/nar/gkz1062.

112. Puente-Santamaria, L., Wasserman, W.W., and Del Peso, L. (2019). TFEA.ChIP: a tool kit for transcription factor binding site enrichment analysis capitalizing on ChIP-seq datasets. Bioinformatics 35, 5339–5340. 10.1093/bioinformatics/btz573.

113. Jing Wang, Y.L., Eric Jaehnig, Zhiao Shi, Quanhu Sheng (2023). Gene Set Analysis Toolkit WebGestaltR. https://github.com/bzhanglab/WebGestaltR.

114. James Li, J.B., Martin Wells (2022). Tools for Tensor Analysis and Decomposition. https://github.com/rikenbit/rTensor.

115. Schindelin, J., Arganda-Carreras, I., Frise, E., Kaynig, V., Longair, M., Pietzsch, T., Preibisch, S., Rueden, C., Saalfeld, S., Schmid, B., et al. (2012). Fiji: an open-source platform for biological-image analysis. Nat Methods 9, 676–682. 10.1038/nmeth.2019.

116. Gray, D.H., Chidgey, A.P., and Boyd, R.L. (2002). Analysis of thymic stromal cell populations using flow cytometry. J Immunol Methods 260 (*1-2*), 15–28. 10.1016/s0022-1759(01)00493-8.

117. Zheng, G.X., Terry, J.M., Belgrader, P., Ryvkin, P., Bent, Z.W., Wilson, R., Ziraldo, S.B., Wheeler, T.D., McDermott, G.P., Zhu, J., et al. (2017). Massively parallel digital transcriptional profiling of single cells. Nat Commun 8, 14049. 10.1038/ncomms14049.

