## Supplemental data for "Coordinated changes in thymic stromal and hematopoietic cells that define the perinatal to juvenile transition"

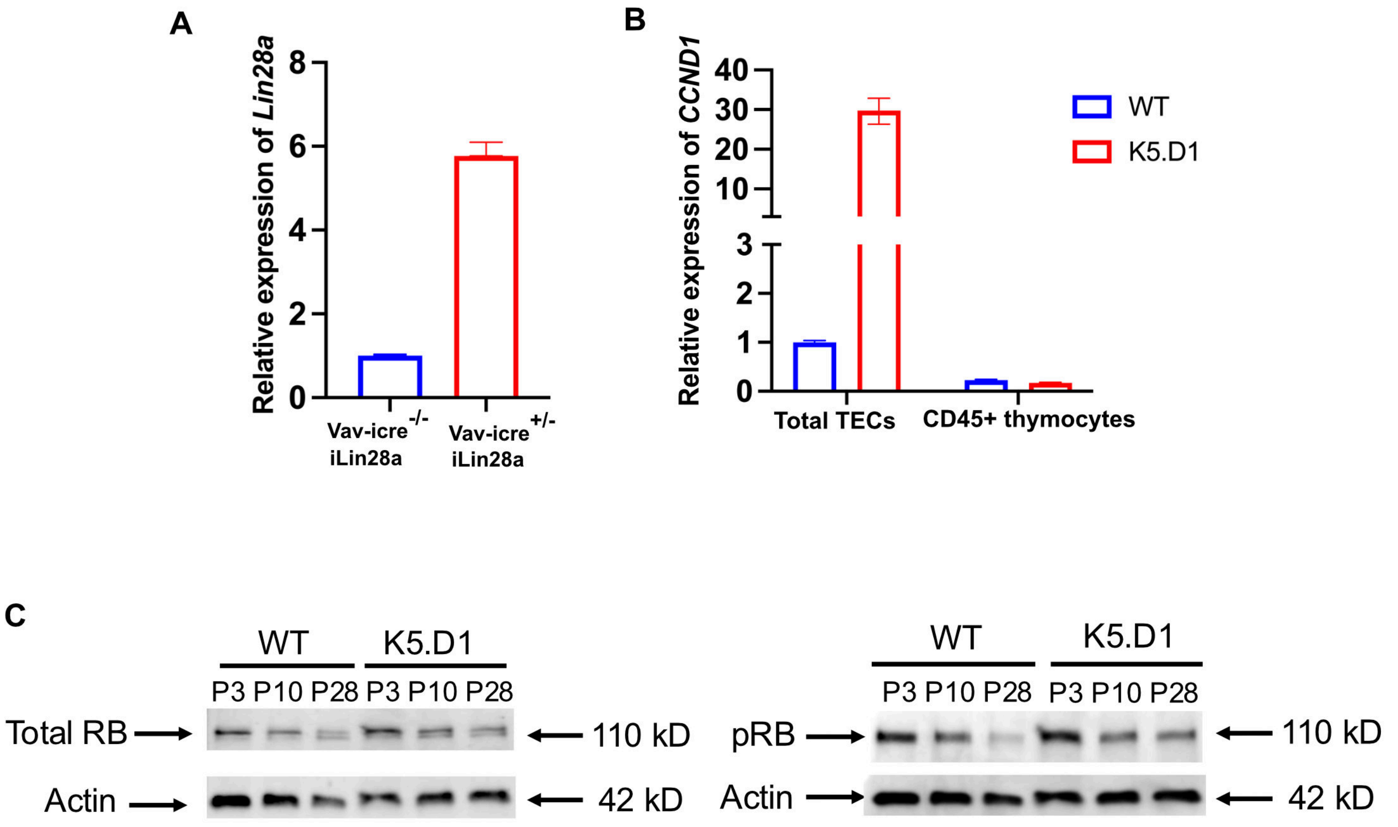


**Figure S1, related to Figure 1: TECs regulate perinatal growth transition.**

(A) RT-qPCR data showing *Lin28a* expression in bone marrow from 8-week-old recipient *Foxn1$^{lacZ}$* mutant mice as previously reported[18]. BM cells ($1x10^7$) from 9-week-old Vav-iCre$^{+/-}$;R26iLin28a and Vav-iCre$^{-/-}$;R26iLin28a mice that were T/B cell depleted were transferred into lethally irradiated 8 week old *Foxn1$^{lacZ}$* mutants (+/Z and ZZ mice). After 21 days, donor BM cells were analyzed in recipient mice. Data are obtained from 3 independent experiments. Statistical analysis was performed using student's t-test *P<0.05, **P<0.01, ***P<0.0001, and ****P< 0.00001. Expression levels were normalized to Vav-iCre$^{-/-}$;R26iLin28a mice.

(B) Representative RT-qPCR data showing relative expression of *CCND1* in total TECs and CD45+ thymocytes from WT vs K5.D1 mice. Expression levels for total TECs and CD45 thymocytes were normalized to WT TEC sample.

(C) Representative western blot images showing total Rb (left) or phosphorylated Rb (right) and actin bands in WT and K5.D1 mice at the indicated ages.

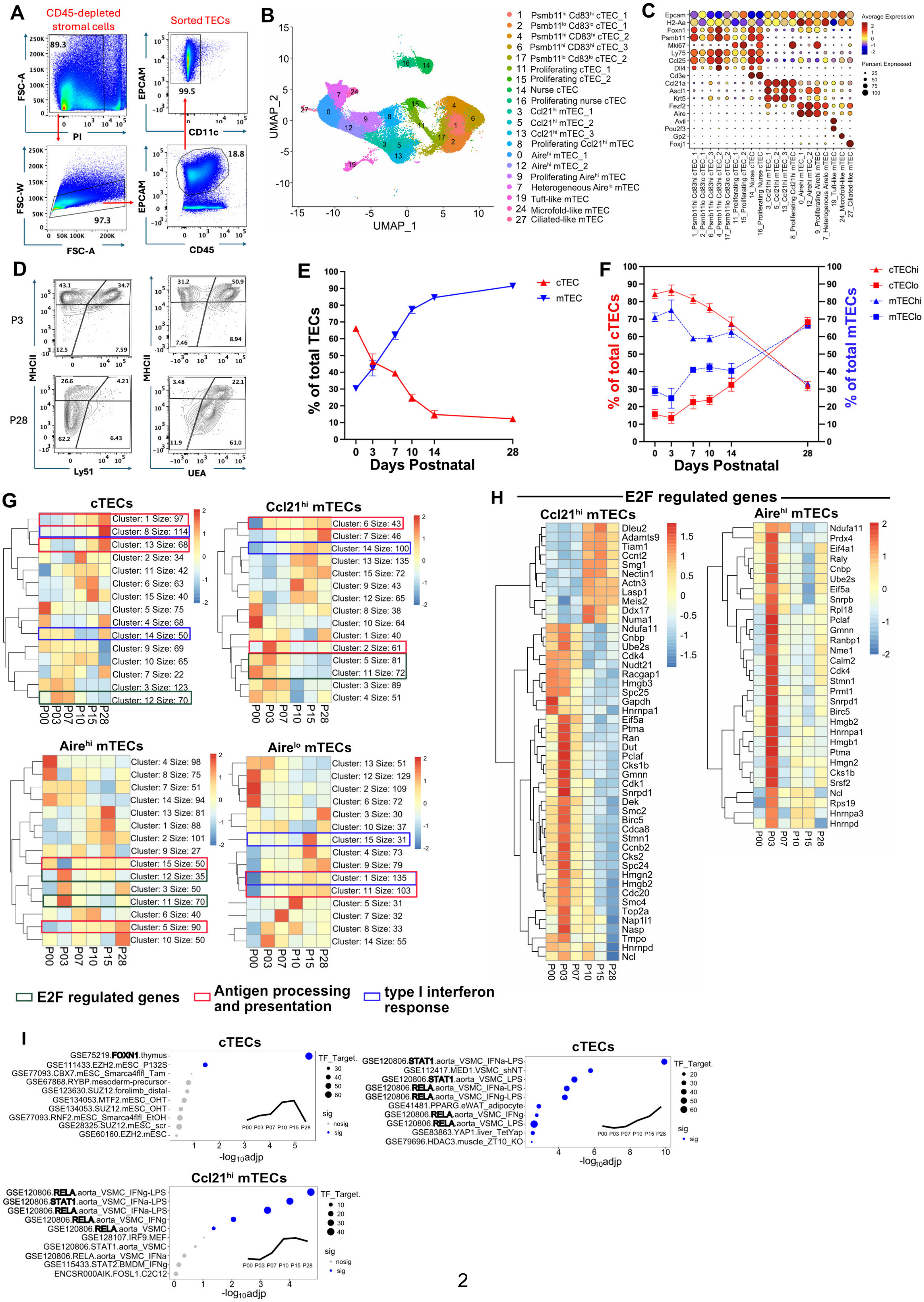
A
CD45-depleted stromal cells
Sorted TECs
FSC-A
PI
89.3
EPCAM
CD11c
99.5
FSC-W
FSC-A
97.3
EPCAM
CD45
18.8
B
UMAP_2
UMAP_1
1 Psmb11hi Cd83hi cTEC_1
2 Psmb11lo Cd83lo cTEC_1
4 Psmb11hi CD83hi cTEC_2
6 Psmb11hi Cd83hi cTEC_3
17 Psmb11lo Cd83lo cTEC_2
11 Proliferating cTEC_1
15 Proliferating cTEC_2
14 Nurse cTEC
16 Proliferating nurse cTEC
3 Ccl21hi mTEC_1
5 Ccl21hi mTEC_2
13 Ccl21hi mTEC_3
8 Proliferating Ccl21hi mTEC
0 Airehi mTEC_1
12 Airehi mTEC_2
9 Proliferating Airehi mTEC
7 Heterogeneous Airelo mTEC
19 Tuft-like mTEC
24 Microfold-like mTEC
27 Ciliated-like mTEC
C
Epcam
H2-Aa
Foxn1
Psmb11
Mki67
Ly75
Ccl25
Dll4
Cd3e
Ccl21a
Ascl1
Krt5
Fezf2
Aire
Avil
Pou2f3
Gp2
Foxj1
Average Expression
Percent Expressed
D
P3
P28
MHCII
Ly51
UEA
E
% of total TECs
Days Postnatal
cTEC
mTEC
F
% of total cTECs
% of total mTECs
Days Postnatal
cTEChi
cTEClo
mTEChi
mTEClo
G
cTECs
Ccl21hi mTECs
Airehi mTECs
Airelo mTECs
P00 P03 P07 P10 P15 P28
E2F regulated genes
Antigen processing and presentation
type I interferon response
H
E2F regulated genes
Ccl21hi mTECs
Airehi mTECs
I
cTECs
TF_Target.
sig
-log10adjp
cTECs
Ccl21hi mTECs
P00 P03 P07 P10 P15 P28

**Figure S2, related to Figure 2: Changes in transcriptional profiles and subset composition of TECs across perinatal to adult transition**

(A) Gating strategy used to isolate $CD45^-$ depleted total TECs for scRNAseq analysis.

(B) UMAP of TEC scRNAseq data from Figure 2A with annotation of higher resolution clusters that further subdivide major TEC subsets.

(C) Dot plot shows average expression of genes used to annotate higher resolution TEC subsets.

(D) Representative flow cytometry plots of $EPCAM^+CD45^-$ TECs showing the frequencies of $cTEC^{hi}$ ($Ly51^+MHCII^{hi}$), $cTEC^{lo}$ ($Ly51^+MHCII^{lo}$), $mTEC^{hi}$ ($UEA^+MHCII^{hi}$) and $mTEC^{lo}$ ($UEA^+MHCII^{lo}$) TECs from P3 and P28 mice.

(E) Quantification of the percentage of total cTECs ($Ly51^+$) and mTECs ($UEA^+$). n=8-26 mice per age group.

(F) Quantification of the frequencies of $cTEC^{hi}$ and $cTEC^{lo}$ subsets within the cTEC compartment (red) and $mTEC^{hi}$ and $mTEC^{lo}$ subsets within the mTEC compartment (blue), from flow cytometry data as in (D).

(G) Heatmaps showing average expression levels of gene modules identified by k-means clustering of scRNAseq data from the indicated TEC subsets. Modules enriched for E2F regulated genes (green boxes), type I IFN regulated genes (blue boxes), and antigen processing and presentation pathways (red boxes) are indicated.

(H) Heatmap showing normalized expression levels of E2F regulated genes, from GSEA of k-means clusters from (G), in $Ccl21^{hi}$ TEC and $Aire^{hi}$ mTEC subsets.

(I) The top 10 most significantly enriched transcription factors regulating a specific time-dependent gene expression pattern (shown in the subgraph) in cTECs (top) and $Ccl21^{hi}$ TECs (bottom). Node size represents the number of overlapping genes between the selected gene set and transcription factor binding targets. Nodes in blue indicate statistically significant enrichment (adjp < 0.05), while gray denotes non-significant enrichment.

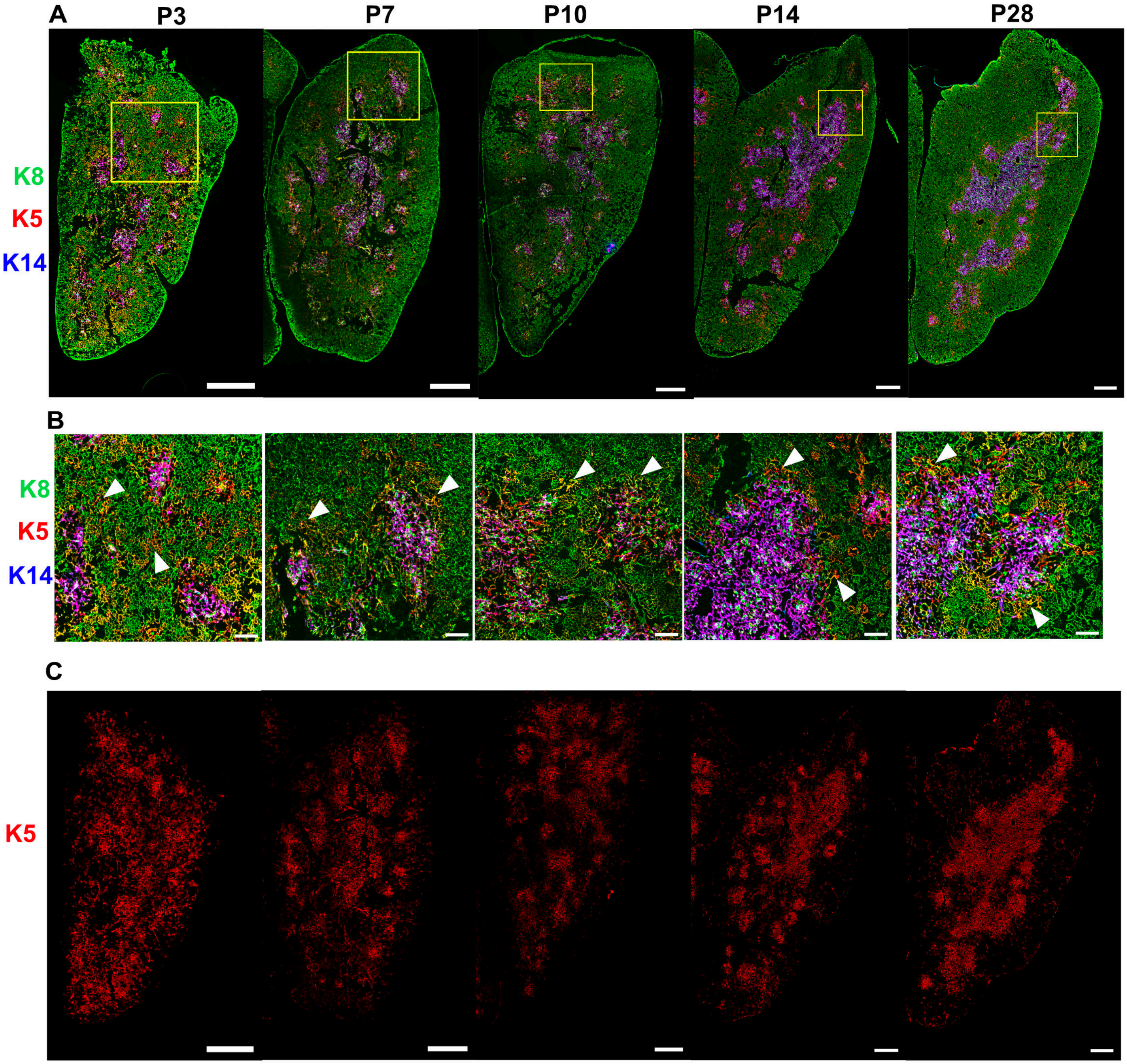


**Figure S3, related to Figure 2: Changes in cortical and medullary organization across perinatal to adult transition**

(A) Representative immunofluorescence images of WT P3, P7, P10, P14 and P28 thymus cryosections. Sections were stained with antibodies to Krt8 (green), Krt5 (red) and Krt14 (blue). Yellow boxes show regions magnified in (B). Scale bars from these 10X images = 500μm. Images are representative of thymus sections from 3 mice.

(B) Magnified images from yellow boxes in (A). White arrows identify K8 and K5 co-staining TECs. Scale bars from these images = 100μm

(C) Single-channel immunofluorescence images from data shown in (A), showing only Krt5 immunostaining. Scale bars from these 10X images = 500μm

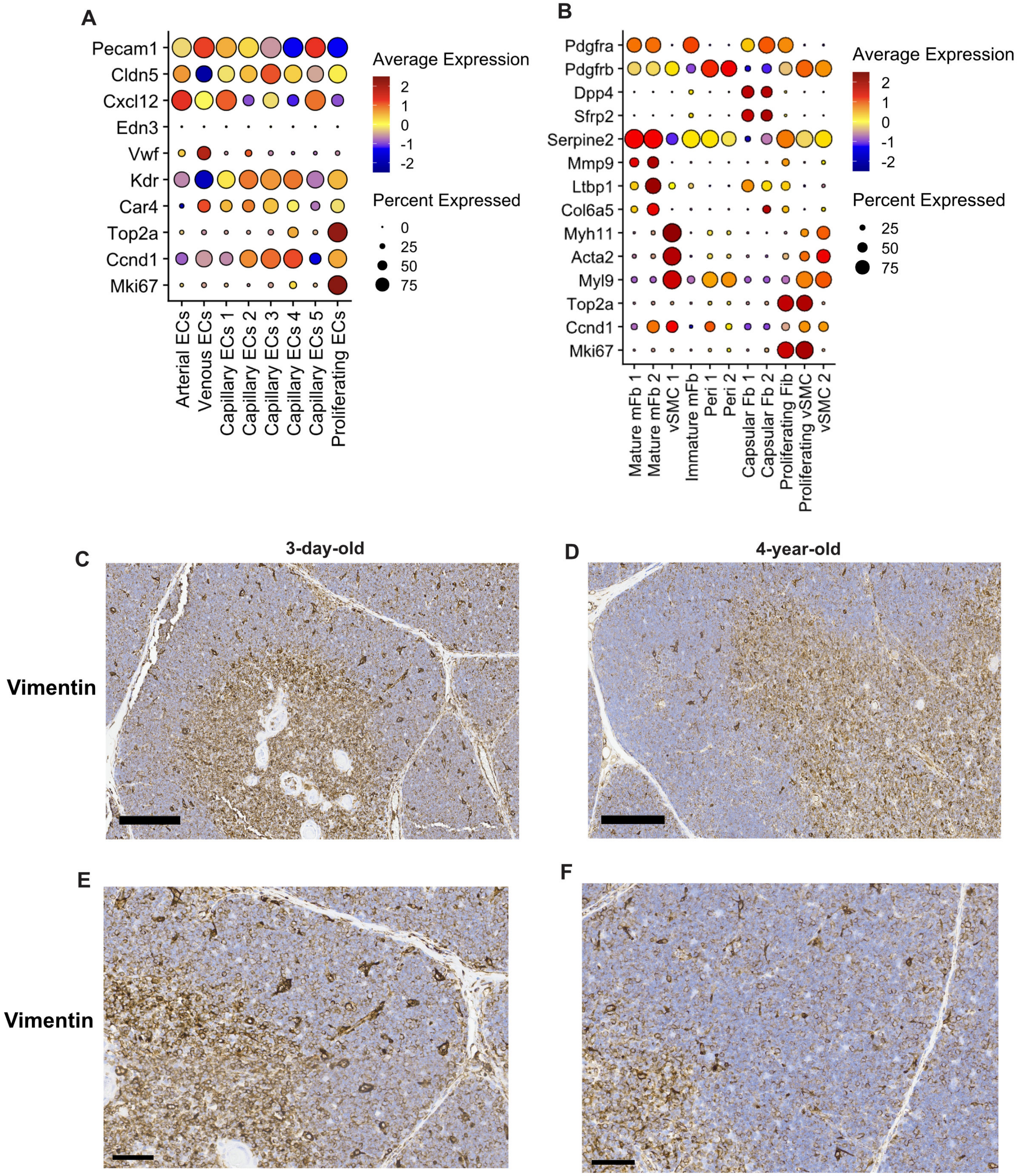


**Figure S4, related to Figure 3: Fibroblast and endothelial cell distribution across perinatal stages.**
(A) dotplot showing genes used to identify subsets endothelial in Figure 3F.
(B) dotplot showing genes used to identify mesenchymal subsets in Figure 3H.
(C, D) IHC of Vimentin+ fibroblasts in 3-day-old & 4-year-old human thymi, respectively. Scale bar = 200μm.
(E, F) Magnified view of Vimentin+ fibroblasts from (C,D). scale bar = 60 μm

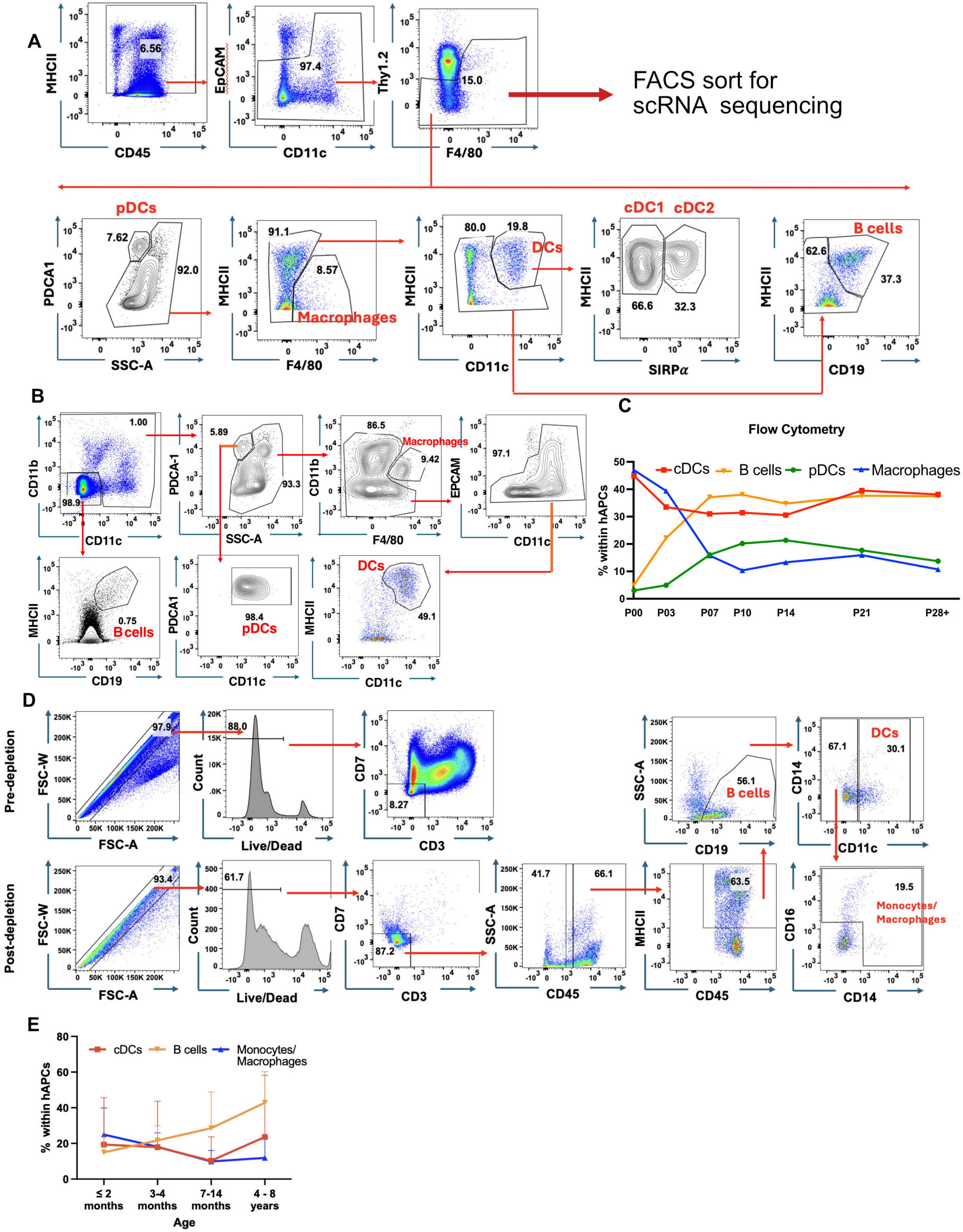
A
FACS sort for scRNA sequencing
pDCs
Macrophages
DCs
cDC1 cDC2
B cells
B
C
Flow Cytometry
cDCs B cells pDCs Macrophages
% within hAPCs
P00 P03 P07 P10 P14 P21 P28+
D
Pre-depletion
Post-depletion
Monocytes/ Macrophages
E
cDCs B cells Monocytes/ Macrophages
% within hAPCs
≤ 2 months 3-4 months 7-14 months 4 - 8 years
Age

**Figure S5, related to Figure 4: scRNAseq and flow cytometry reveal cellular changes in mouse and human hAPC composition over the transition**

(A) Representative gating strategy used to FACS sort hAPCs for scRNAseq analysis. The second row of flow cytometry plots were analyzed to validate that the gating strategy included all major hAPC cell types. Cells are pre-gated on live single cells.

(B) Representative flow cytometry plots display the gating strategy used to quantify thymic hAPC subsets from mouse thymus across the transition in Figure 4C. Cells are pre-gated on live single cells.

(C) Quantification of cDCs, B cells, pDCs and macrophages, as a frequency of total hAPCs, based on flow cytometry data as in (B).

(D) Representative flow cytometry plots display the gating strategy used to identify thymic hAPC subsets from human thymus. Single-cell suspensions generated from thymus tissue were depleted of CD3+ and CD7+ cells using magnetic beads and then stained with fluorescent antibodies to identify the labeled hAPC subsets. (Top panel) Sample prior to depletion showing singlet gates, viability and CD3/CD7 staining. (Bottom panel) The same sample after depletion. The live $CD3^-CD7^-CD45^+MHCII^+$ APC populations were characterized as B cells ($CD19^+$), DCs ($CD19^-CD11c^+$), or monocytes ($CD19^-CD11c^-$ then $CD14^+$, $CD14^+CD16^+$ or $CD16^+$).

(E) Quantification of cDCs, B cells and monocytes as a frequency of hAPCs identified by flow cytometry in human samples across the perinatal to adult transition, as per the gating strategy in (S5D).

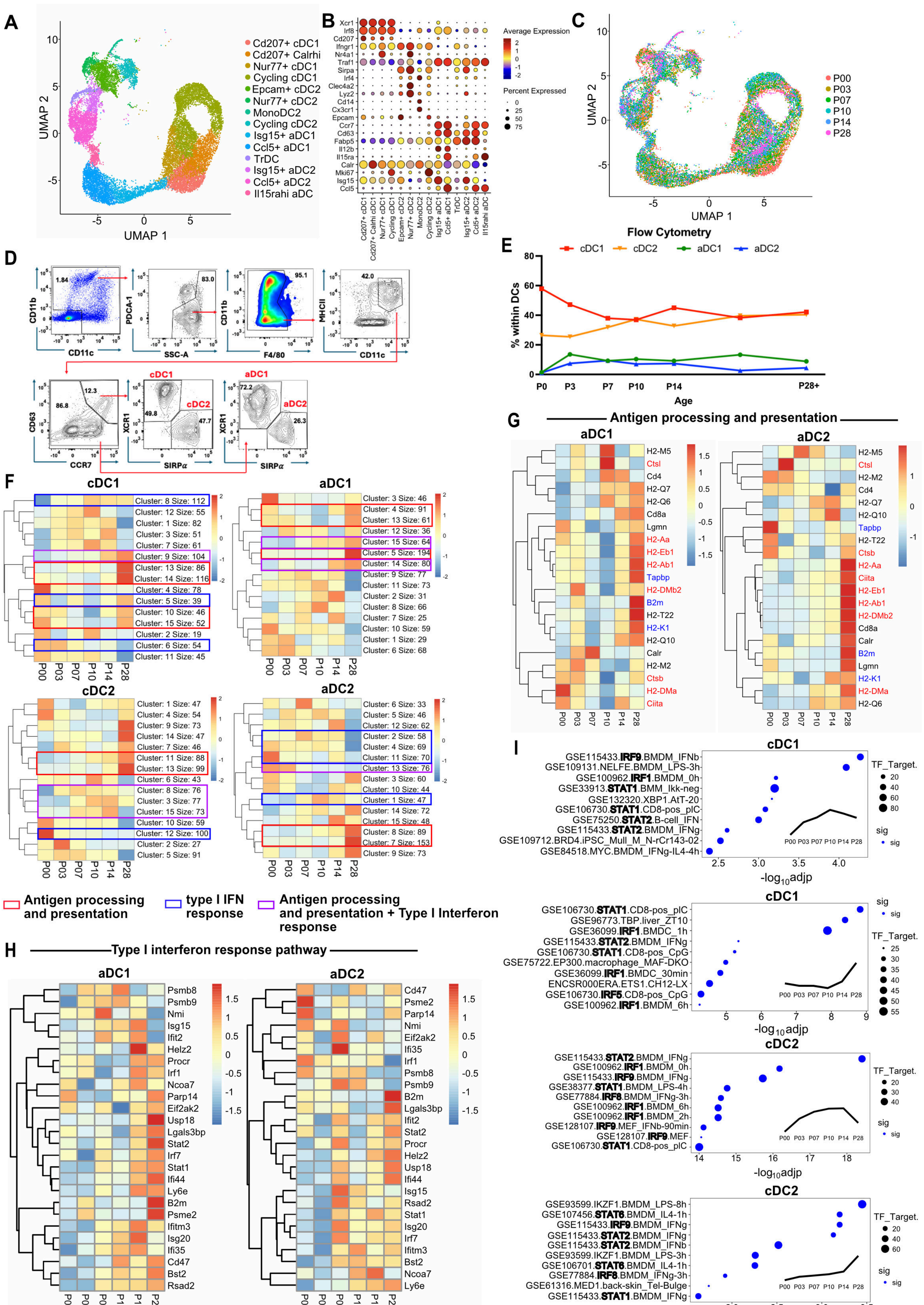
A
Cd207+ cDC1
Cd207+ Calrhi
Nur77+ cDC1
Cycling cDC1
Epcam+ cDC2
Nur77+ cDC2
MonoDC2
Cycling cDC2
Isg15+ aDC1
Ccl5+ aDC1
TrDC
Isg15+ aDC2
Ccl5+ aDC2
Il15rahi aDC
UMAP 1
UMAP 2
B
Average Expression
Percent Expressed
Xcr1
Irf8
Cd207
Ifngr1
Nr4a1
Traf1
Sirpa
Irf4
Clec4a2
Lyz2
Cd14
Cx3cr1
Epcam
Ccr7
Cd63
Fabp5
Il12b
Il15ra
Calr
Mki67
Isg15
Ccl5
C
P00
P03
P07
P10
P14
P28
D
CD11b
CD11c
PDCA-1
SSC-A
F4/80
MHCII
CD63
CCR7
XCR1
SIRPα
cDC1
cDC2
aDC1
aDC2
E
Flow Cytometry
cDC1
cDC2
aDC1
aDC2
% within DCs
Age
P0
P3
P7
P10
P14
P28+
F
cDC1
aDC1
cDC2
aDC2
Antigen processing and presentation
type I IFN response
Antigen processing and presentation + Type I Interferon response
G
Antigen processing and presentation
aDC1
aDC2
H
Type I interferon response pathway
aDC1
aDC2
I
cDC1
cDC2
TF_Target.
sig
-log10adjp
GSE115433.IRF9.BMDM_IFNb
GSE109131.NELFE.BMDM_LPS-3h
GSE100962.IRF1.BMDM_0h
GSE33913.STAT1.BMM_Ikk-neg
GSE132320.XBP1.AtT-20
GSE106730.STAT1.CD8-pos_pIC
GSE75250.STAT2.B-cell_IFN
GSE115433.STAT2.BMDM_IFNg
GSE109712.BRD4.iPSC_Mull_M_N-rCr143-02
GSE84518.MYC.BMDM_IFNg-IL4-4h
GSE106730.STAT1.CD8-pos_pIC
GSE96773.TBP.liver_ZT10
GSE36099.IRF1.BMDC_1h
GSE115433.STAT2.BMDM_IFNg
GSE106730.STAT1.CD8-pos_CpG
GSE75722.EP300.macrophage_MAF-DKO
GSE36099.IRF1.BMDC_30min
ENCSR000ERA.ETS1.CH12-LX
GSE106730.IRF5.CD8-pos_CpG
GSE100962.IRF1.BMDM_6h
GSE115433.STAT2.BMDM_IFNg
GSE100962.IRF1.BMDM_0h
GSE115433.IRF9.BMDM_IFNg
GSE38377.STAT1.BMDM_LPS-4h
GSE77884.IRF8.BMDM_IFNg-3h
GSE100962.IRF1.BMDM_6h
GSE100962.IRF1.BMDM_2h
GSE128107.IRF9.MEF_IFNb-90min
GSE128107.IRF9.MEF
GSE106730.STAT1.CD8-pos_pIC
GSE93599.IKZF1.BMDM_LPS-8h
GSE107456.STAT6.BMDM_IL4-1h
GSE115433.IRF9.BMDM_IFNg
GSE115433.STAT2.BMDM_IFNg
GSE115433.STAT2.BMDM_IFNb
GSE93599.IKZF1.BMDM_LPS-3h
GSE106701.STAT6.BMDM_IL4-1h
GSE77884.IRF8.BMDM_IFNg-3h
GSE61316.MED1.back-skin_Tel-Bulge
GSE115433.STAT1.BMDM_IFNg

**Figure S6, related to Figure 4: scRNAseq reveals changes in DC composition and transcriptional signatures reflecting increased Type I IFN signaling over the transition**

(A) UMAP of cDC scRNAseq data from Figure 4E with annotation of higher resolution clusters that further subdivide major cDC subsets. (n=2 per age).

(B) Dot plot shows average normalized expression values of genes used to annotate higher resolution cDC subsets in A.

(C) UMAP from (A), color-encoded by age.

(D) Flow cytometry plots showing the representative gating strategy used to identify thymic cDC subsets in mice. Cells are pre-gated on live single cells.

(E) Quantification of cDC1s, cDC2s, aDC1s, and aDC2s as a frequency of total cDCs defined by flow cytometry, as shown in (D), in mouse thymus across the perinatal to adult transition.

(F) Heatmaps showing average expression levels of gene modules identified by k-means clustering of scRNAseq data from the indicated cDC subsets. Modules enriched for type I IFN regulated genes (blue boxes), antigen processing and presentation pathways (red boxes), or both (purple boxes) are indicated.

G) Heatmaps showing normalized expression of genes associated with the antigen processing and presentation pathway, from GSEA of k-means clusters (Figure S6F), in aDC1 and aDC2 subsets at the indicated ages across the perinatal to juvenile transition. Genes highlighted indicate MHCI genes (blue) and MHCII genes (red).

(H) Heatmaps showing normalized expression levels of Type I interferon-regulated genes, from GSEA of k-means clusters from (Figure S6F), in aDC1 and aDC2 subsets at the indicated ages.

(I) The top 10 most significantly enriched transcription factors regulating a specific time-dependent gene expression pattern (shown in the subgraph) in cDC1s and cDC2s. Node size represents the number of overlapping genes between the selected gene set and transcription factor binding targets. Nodes in blue indicate statistically significant enrichment (adjp < 0.05), while gray denotes non-significant enrichment.

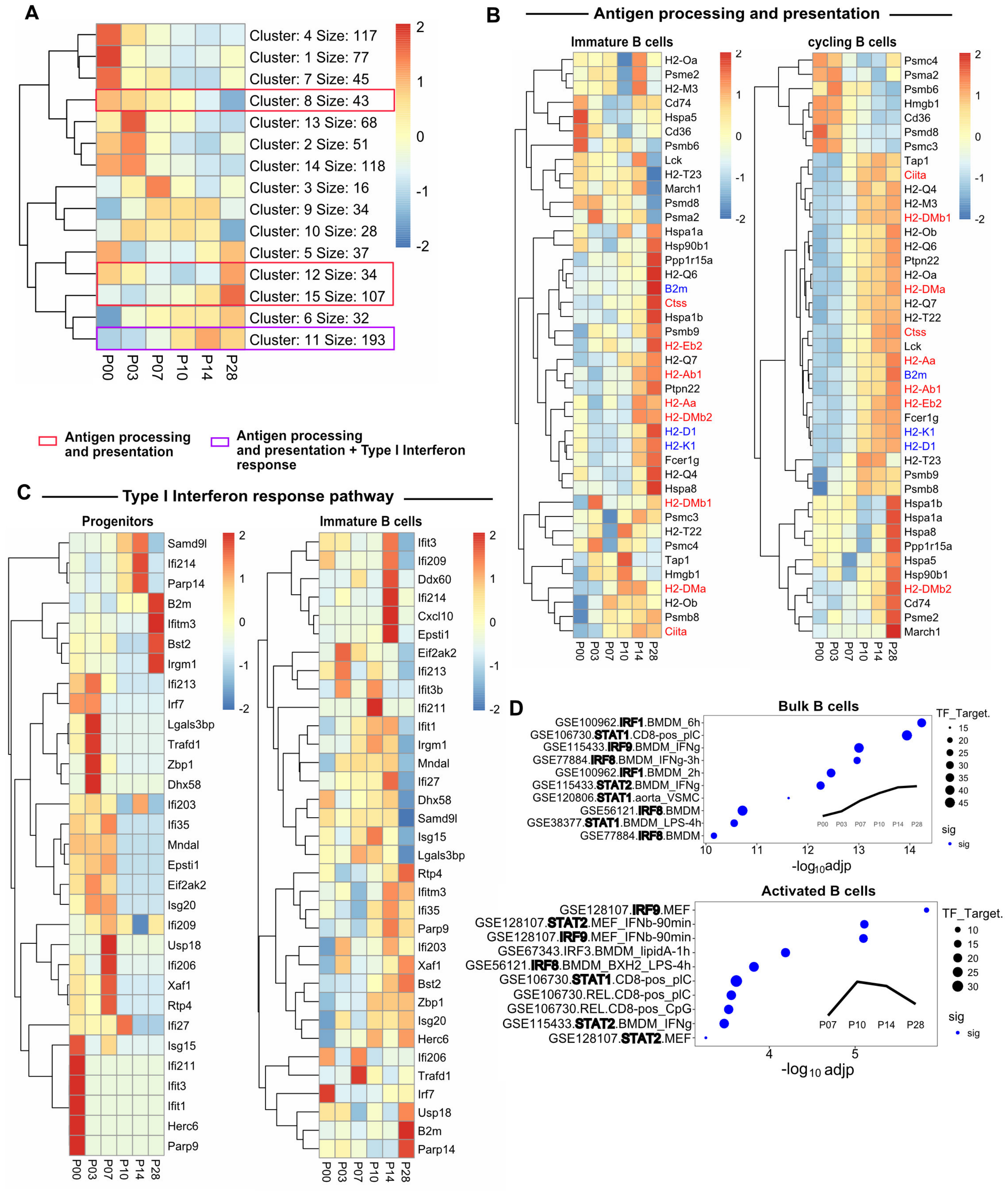


**Figure S7, related to Figure 5: scRNAseq reveals changes in transcriptional signatures in B cells, reflecting increased Type I IFN signaling and altered antigen processing and presentation over the transition.**

(A) Heatmaps showing average expression levels of gene modules identified by k-means clustering of scRNAseq data from thymic B cells. Modules enriched in antigen processing and presentation pathways alone (red boxes) or together with type I IFN regulated genes (purple boxes) are indicated.
(B) Heatmaps showing normalized expression values of genes associated with the Antigen Processing and Presentation pathway, from GSEA of k-means clusters (Figure S7A), in immature and cycling B cell subsets at the indicated ages. Genes highlighted indicate MHCI genes (blue) and MHCII genes (red).
(C) Heatmap showing normalized expression values of genes associated with the Type I IFN Response pathway, from GSEA of k-means clusters (Figure S7A) in progenitor and Immature B cell subsets at the indicated ages.
(D) The top 10 most significantly enriched transcription factors regulating a specific time-dependent gene expression pattern (shown in the subgraph) in B cells activated B cells. Node size represents the number of overlapping genes between the selected gene set and transcription factor binding targets. Nodes in blue indicate statistically significant enrichment (adjp < 0.05), while gray denotes non-significant enrichment.

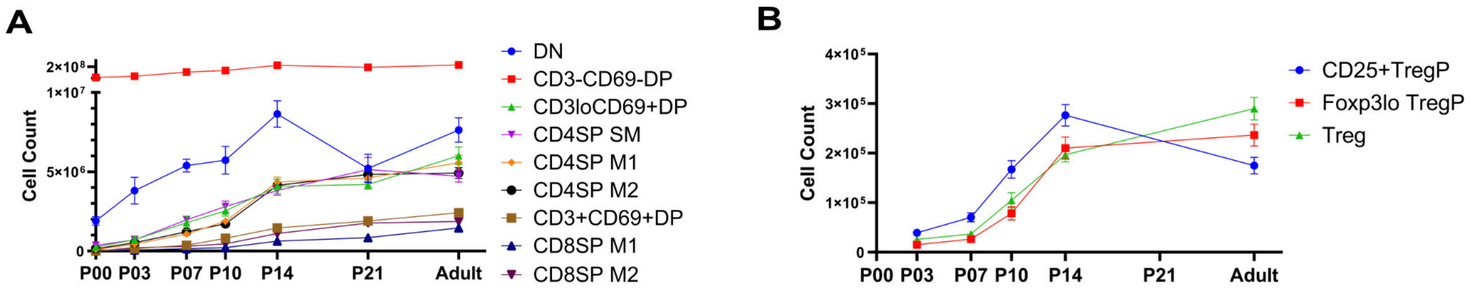


**Figure S8, related to Figure 6: The cellularity of conventional thymocyte subsets and Treg is altered over the perinatal to juvenile transition.**

(A) Quantification of thymocyte subsets at the indicated ages. Data are pooled from at least 3 independent experiments per age group (n=8-47 mice per age group). Symbols with error bars represent mean ±SEM.
(B). Quantification of thymic CD25+TregP, Foxp3lo TregP, and Tregs. Data are pooled from at least 3 independent experiments per age group (n=9-30 mice per age group). Symbols with error bars represent mean±SEM.
